# PhageTAILor leverages machine learning for phage tail-like elements detection and classification in plant-associated bacteria

**DOI:** 10.64898/2026.08.24.746745

**Authors:** Heejung Cho, Socheata Hour, Simon Roux, Clement Coclet, Oluwamayowa Amusat, Vivek Mutalik, Alexey Kazakov, Asaf Levy, Nimrod Nachmias, Lorenzo Aureli, Tyrome Sweet, Axel Visel, Ruben Michael Ceballos, Jonelle TR Basso

## Abstract

Phage tail-like elements (PTEs) — tailocins, bacterial type VI secretion systems (T6SS), and extracellular contractile injection systems (eCIS) — are contractile nanomachines that bacteria use to kill their neighbors and compete within their micro-ecosystems. PTEs help shape microbial community composition. Most PTE detection tools only detect a single PTE class. Moreover, most tailocin detection methods are largely restricted to *Pseudomonas*, leaving a key part of tailocin diversity uncharacterized. In this work, we present PhageTAILor (https://github.com/hjcho-bio/PhageTAILor), an integrative and fully automated pipeline that detects and classifies prophages and 3 PTE classes from bacterial genomes. PhageTAILor combines a 6-detector homology-based candidate search (geNomad, tail-gene, PHROGs-tail, SecReT6, eCIStem, and a divergence-tolerant tail-HMM detector) with a LightGBM classifier comprising 1 multiclass and 3 binary heads, trained on 6,501 bacterial genomes carrying 13,082 prophages and PTEs. A phylogeny-free feature matrix used in our model keeps predictions reproducible between model construction and user inference. PhageTAILor performs strongly at the genome level and generalizes beyond its *Pseudomonas*-rich training set. On a 76-strain cross-clade benchmark, PhageTAILor detected tailocins at F1 = 0.955. Furthermore, it identified 12 of 13 experimentally validated tailocins spanning five genera versus 2 of 13 for a *Pseudomonas*-restricted tool TattleTail. PhageTAILor also demonstrated sensitivity equivalent to viral detection tool geNomad while avoiding its higher false-positive rate. Applied to 7,925 plant- and soil-associated bacterial isolates, PhageTAILor showed that prophages in the phyllosphere and tailocins in plant-associated bacteria, whereas eCIS are enriched in soil. PhageTAILor is distributed as an open-source, modular pipeline with a command-line interface.

**IMPORTANCE:** Bacteria produce phage tail-like elements — including tailocins, type VI secretion systems, and extracellular contractile injection systems — to kill competitors and inject toxins, processes that help determine interbacterial and host-bacterial interactions as well as microbial community structures. Identifying these elements across broad bacterial phylogeny has been limited because current tools target only a single element class or designed to work almost exclusively on *Pseudomonas* strains. PhageTAILor addresses this by detecting and classifying prophages and 3 PTE classes from any bacterial genome and, critically, generalizing to bacterial lineages far from its training data. In addition to offering an accurate and automated detection framework, applying PhageTAILor across large datasets uncovers broad ecological structuring in how these competition systems are distributed. PhageTAILor provides a scalable framework for investigating interbacterial competition, microbiome assembly, and phage-derived antimicrobials.

## INTRODUCTION

PTEs are contractile or syringe-like nanomachines of bacteria that are repurposed from phage tail ancestors to mediate interbacterial competition and host-bacterial interactions by delivering toxic effectors or disrupting membrane polarity. In this work, we use the term PTEs to include tailocins, T6SSs, and eCISs. While PTEs broadly influence microbial community structure, host interactions, and nutrient cycling, their heterogeneous and frequently fragmented genomic signatures make large-scale automated identification challenging (1–4).

Computational tools have been developed that use characteristic marker sets to detect prophages and each PTE class, enabling substantial progress to understand individual PTE types (2–5). However, there is a need for an integrated, unified platform that can detect multiple PTE classes by assimilating diverse evidence sources, thereby expanding our ability to discover non-canonical architectures, cross-clade variants, and systems in even under-sampled taxa. Prophages have been studied far more extensively than PTEs. Indeed, a myriad of viral detection tools such as geNomad, PHASTEST, and VirSorter2 have been used to detect phages and prophages in various host organisms in microbiome research, therapeutics and clinical studies, biotechnological engineering and many other fields (5–8). Computational predictions for prophages and tailocins typically rely on virus-centric detectors or PHROG-based tail annotations (9–11). To date only one study describes a publicly available tailocin detector tool called TattleTail (2). TattleTail uses conserved pyocin gene cluster markers and uses the absence of many canonical phage features that are not found in tailocins to detect tailocins, exclusively in *Pseudomonas.* As the most widely studied PTE class, T6SS characterization has advanced rapidly through curated databases like SecReT6 and detection frameworks such as T346Hunter and MacSyFinder (4, 12, 13). Applying these tools has expanded known T6SS distributions across diverse taxa, including Bacteroidota, Francisella, and Pseudomonadota, and are divided into subtypes i, ii, and iii with distinct architectures and taxonomic ranges (14–16). In contrast, extracellular contractile injection systems (eCIS)—though biologically important and increasingly recognized in environmental lineages—remain sparsely annotated and are frequently missed by phage-centric pipelines. To address this, tools like eCIStem have begun mapping these loci using target marker profiles (3, 17). Recent work by Nachmias et al. (18) exemplifies the application of the eCIStem database in detecting eCIS-binding domains to gain deeper insights in diversity, microbial interactions, functional adaptation and evolutionary changes in 1,069 bacteria and archaea. Although these previous works provide invaluable insights to expand our ability to detect PTEs, comparative analysis of PTE distribution across niches, hosts, and phylogenetic lineages still lacks a unified, scalable, and well-calibrated detection framework.

Tailocins are contact-dependent, narrow-spectrum weapons that kill competitors (19). The rhizosphere and endosphere are dense, spatially structured, and nutrient-contested habitats colonized by microbes competing for the same root exudates in the soil interface and entry sites for internal colonization (20, 21). Root-driven nutrients place high selective pressure on diverse soil bacteria in these environments (22, 23). Contact-dependent weapons like tailocins may play essential roles in such structured, high-density settings where the producer can kill neighbors locally and monopolizes the colonization site. In contrast, in well-mixed dilute environments, diffusible bacteriocins or no weapon may be favored.

The type VI secretion system (T6SS) is a contact-dependent contractile cellular machinery that Gram-negative bacteria use to inject toxic effectors directly into adjacent cells. It is an inverted, cell-envelope-anchored relative of the contractile phage tail, in which contraction of an outer sheath drives a spike-tipped inner tube across the envelope and into the target(14, 24). Through a broad collection of antibacterial or anti-eukaryotic effectors and their cognate immunity proteins, T6SSs mediate interbacterial competition, host colonization, and niche establishment (15, 25).

eCIS are secreted injection nanomachines whose effectors are typically delivered to eukaryotic target cells such as amoebae, nematodes, and insects (e.g. *Photorhabdus* PVC, entomopathogen/symbiont systems; (18)). Canonical eCIS producers are soil, sediment and animal-associated taxa, including Actinobacteria, Bacteroidota, and entomopathogenic Pseudomonadota (3).

Here, we introduce PhageTAILor, an end-to-end machine-learning pipeline that predicts prophages and three classes of PTEs (tailocins, T6SSs, and eCISs) directly from bacterial genomes by unifying multiple, complementary sequence-evidence sources in a single model. Here, we detail the pipeline architecture and its core outputs, evaluate performance across four benchmark tiers of increasing stringency, compare accuracy against established prophage- and tailocin-detection tools, and demonstrate its utility across 7,925 plant- and soil-associated genomes.

## RESULTS

### Overview of PhageTAILor

PhageTAILor predicts prophages and 3 PTE classes (tailocins, T6SS, and eCIS) directly from bacterial assemblies by unifying various sequence evidence in a single model (Figure 1A). Starting from raw genomic sequence, the pipeline predicts genes and executes six complementary detection modules: geNomad (for proviruses), MMseqs2 searches against curated tail gene databases (PHROGS, SecReT6, and eCIStem), and profile searches against a tail-protein HMM library. Because PTEs are encoded by contiguous gene clusters rather than single genes, each detector aggregates isolated single-gene hits into candidate regions based on scaffold proximity (Figure 1B). Candidates are then represented as 74-feature vectors and evaluated using a LightGBM model (26) containing a five-way multiclass head alongside three class-specific binary heads, with severe class imbalance addressed via per-class loss weighting.

**Figure 1.**
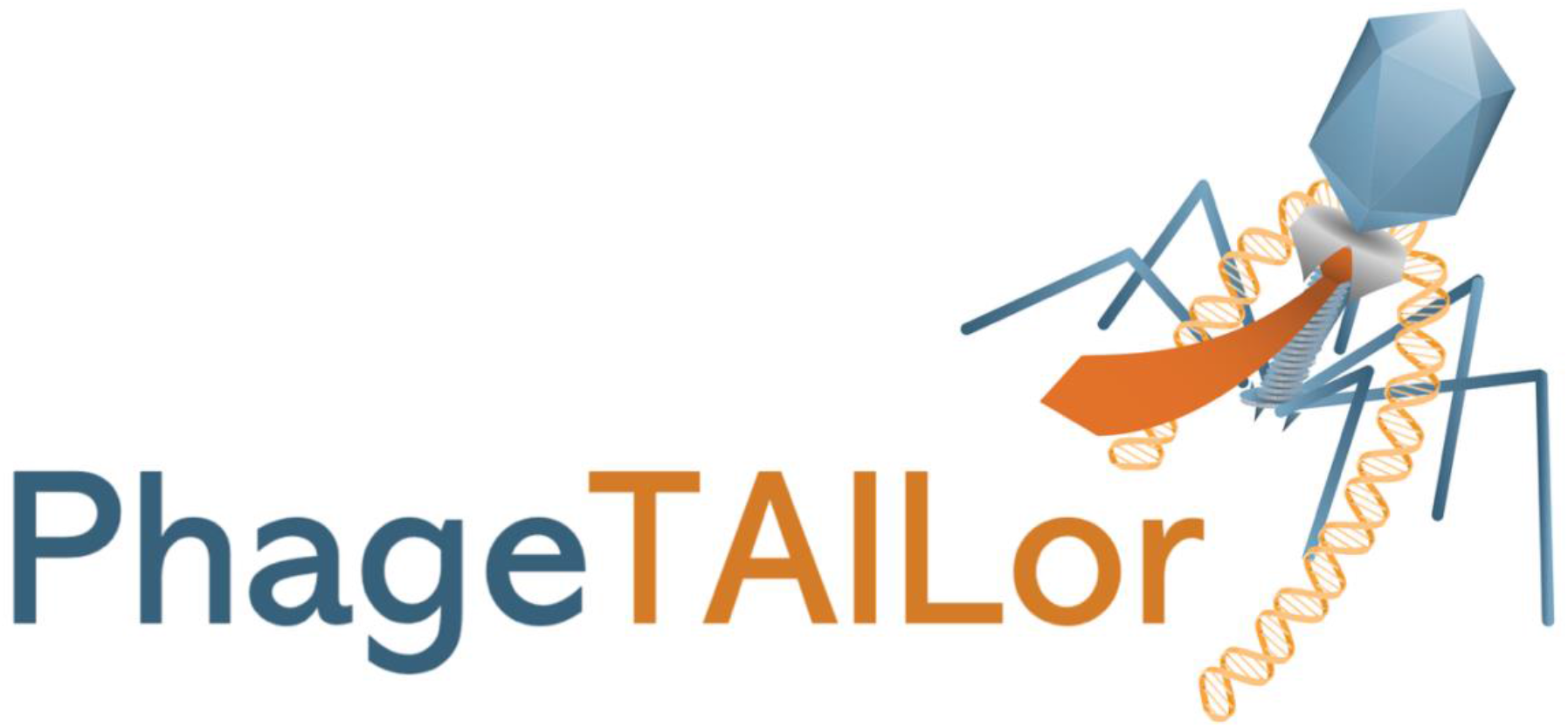
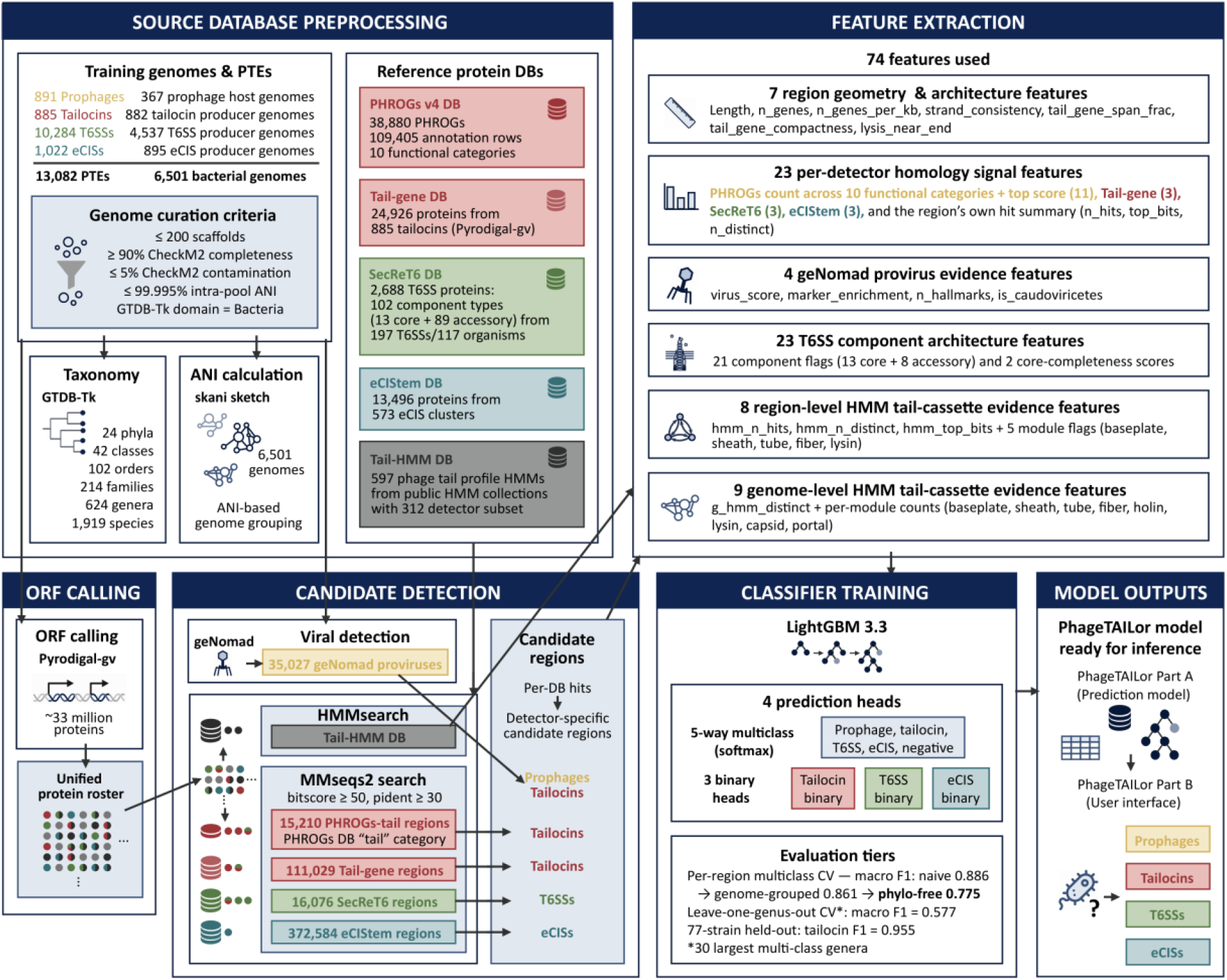
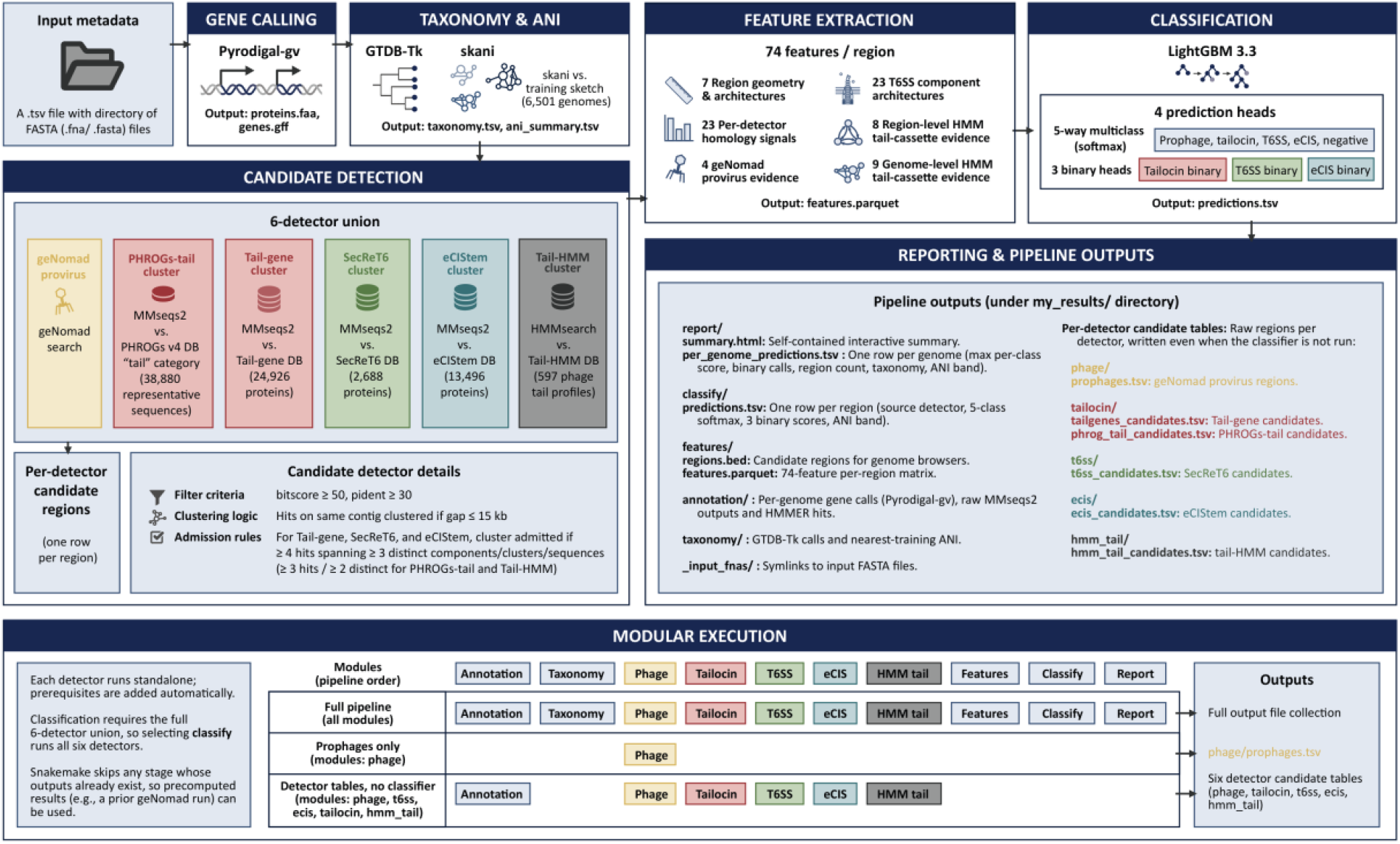
**(A)** PhageTAILor logo. **(B)** The PhageTAILor model-construction workflow (Part A). A curated set of 6,501 quality-controlled bacterial genomes carrying 13,082 PTEs and prophages (891 prophages, 885 tailocins, 10,284 T6SS, 1,022 eCIS) is annotated once with Pyrodigal-gv and searched against five reference databases. Candidate detection by geNomad and four MMseqs2 detectors yields 549,926 candidate regions, while the tail-HMM search adds evidence to feature extraction. Each region becomes a 74-feature vector (region geometry 7, per-detector homology 23, geNomad evidence 4, T6SS architecture 23, and region- and genome-level HMM evidence 8 and 9) that trains a four-head LightGBM classifier (one five-way multiclass head plus tailocin, T6SS, and eCIS binary heads). Across evaluation tiers, per-region CV macro F1 declines from 0.886 to 0.861 to 0.775 (random, genome-grouped, and phylogeny-free), leave-one-genus-out macro F1 is 0.577, and held-out 76-strain tailocin F1 is 0.955. **(C)** PhageTAILor user-inference workflow (Part B). The frozen Part A model is applied to user genomes supplied as a sample file. Pyrodigal-gv calls genes; GTDB-Tk and skani assign taxonomy and the ANI confidence band; and a 6-detector union (geNomad provirus; tail-gene and PHROGs-tail; SecReT6; eCIStem; Tail-HMM) nominates candidate regions under shared filter and clustering rules. Each region becomes a 74-feature vector scored by the four-head LightGBM classifier (one five-way multiclass head plus tailocin, T6SS, and eCIS binary heads). Outputs are per_genome.tsv, predictions.tsv, regions.bed, report.html, and a raw candidate table per detector. The lower panel shows the workflow’s modularity: a module setting selects stages with prerequisites resolved automatically and existing outputs reused, detectors run standalone, and classification requires the full 6-detector union. Example modes are the full pipeline, prophages only, and detector tables without the classifier.

The choice of which head reports each class reflects how the heads behave. Because a 5-way softmax enforces a sum-to-one probability constraint, the presence of another strong PTE type can artificially dilute a genome’s tailocin score. Tailocin and T6SS predictions are therefore drawn from dedicated binary heads, which assess presence or absence independently and remain robust to co-occurring elements. Conversely, eCIS and prophage predictions are sourced from the multiclass head. Under severe class imbalance, the eCIS binary head tends to saturate, frequently overcalling negative genomes that the multiclass model correctly scores near zero. A genome is classified as positive for a given class if the maximum score across its candidate regions—derived from the corresponding model head—exceeds 0.5. To quantify domain shift, each prediction includes an ANI-based confidence band indicating the query genome’s distance from the training set.

The feature set intentionally omits phylogenetic placement variables—specifically label-encoded genus, family, order, and nearest-training ANI. These features cannot be reliably generated during inference and tend to artificially inflate in-distribution accuracy through taxonomy memorization rather than sequence feature learning. Omitting them ensures that our reported benchmarks reliably reflect performance on user-submitted genomes. The model was trained on a curated corpus of 6,501 high-quality genomes containing 13,082 literature- or reference-annotated intervals (10,284 T6SS, 1,022 eCIS, 891 prophages, and 885 tailocins; Figure 5).

### Pipeline outputs

A single command (phagetailor run samples.tsv --out my_results/ --cores 8) executes the complete pipeline and organizes all outputs into a standardized directory structure (Figure 1C). Primary deliverables include a per-region prediction table with all four head probabilities and confidence bands, a per-genome summary that max-pools region-level calls alongside GTDB-Tk taxonomy, a BED file of candidate regions for genomic visualization, and an interactive, browser-readable HTML report. Additionally, each detector outputs an independent candidate table, keeping raw search evidence fully accessible even when running detectors without the machine-learning classifier. For a typical 5-Mb bacterial genome using 8 CPU cores, the pipeline completes in ∼3 minutes, with runtime primarily driven by the geNomad marker search.

### Model evaluation

Detection is evaluated at the genome level and reported across four tiers of increasing stringency, preceded by a per-region leakage check. We first quantified the impact of data leakage on the per-region multiclass cross validation by comparing 3 settings (Figure S1). A five-fold cross validation that split candidate regions at random, with the five phylogenetic placement features included, gave a macro F1 of 0.886. Switching to genome-grouped folds, so that no genome contributed regions to both the training and test sides, lowered this only to 0.861, meaning within genome leakage cost just 0.025 macro F1. Removing the five phylogenetic features then lowered the macro F1 to 0.775 in the deployed phylo-free model, with per class F1 of 0.99 for negative, 0.35 for prophage, 0.84 for tailocin, 0.85 for T6SS, and 0.84 for eCIS. The additional 0.086 of macro F1 that the phylogenetic features accounted for therefore reflects genus-level memorization, and per class F1 remained above 0.84 for every class except prophage without them. We excluded these features because they cannot be replicated at inference time, as a user-provided genome set will rarely reproduce the phylogenetic features present in the training data. All metrics reported below reflect genome-grouped folds evaluated on the phylo-free model.

#### Tier 1. In-distribution genome-level cross-validation

For bacterial taxa present in the training set, T6SS prediction achieved an AUPRC of 0.901 with an F1 score of 0.92 (Figure 2). Tailocin and eCIS predictions yielded high recall (∼1.0); however, precision could not be accurately calculated from this split. Unlabeled genomes in the dataset likely contain an unknown fraction of true positives—particularly tailocins, which are abundant in the *Pseudomonas*-rich training set—thereby artificially depressing precision. To account for this, precision was evaluated using the clean-negative panel in Tier 4. Table 1 lists the production model’s per-class AUPRC across all four classes.

**Figure 2.**
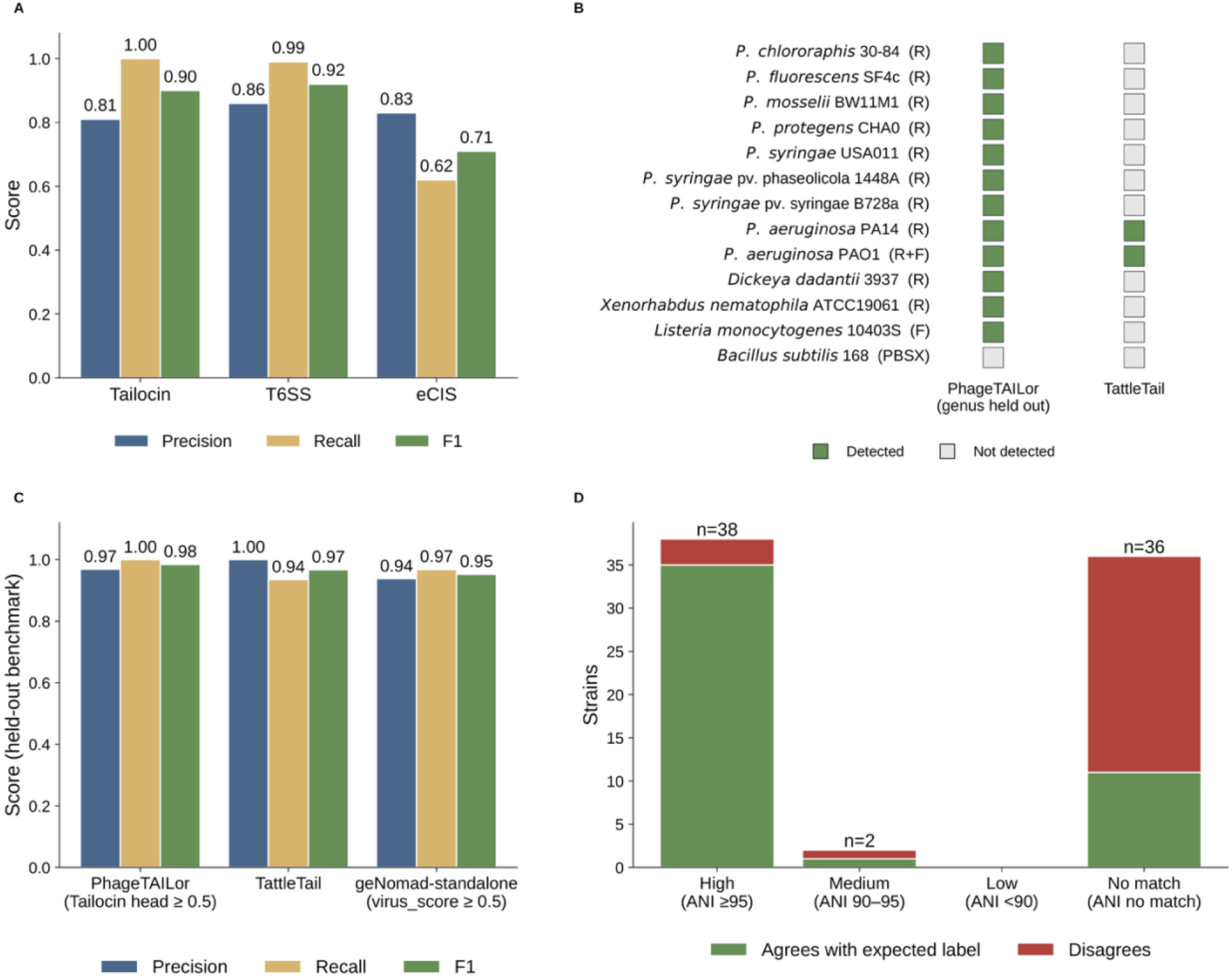
PhageTAILor model performance, cross-genus generalization, comparison with other models, and confidence calibration. **(A)** Genome-level performance of the PhageTAILor model, shown as precision, recall, and F1 for tailocin, T6SS, and eCIS. A genome is called positive for a prophage or PTE class when that class’s head score, taken as the maximum across the genome’s candidate regions, reaches 0.5. Tailocin and T6SS use their binary heads, and eCIS uses the multiclass head. Because no single held-out panel contains enough positives of every class, each class is scored on the most rigorous data available for it. Tailocin precision is taken from the clean-negative panel and tailocin recall from held-out producers, T6SS is scored by genome-grouped cross-validation, and eCIS is scored on the held-out curated benchmark. Prophage is not shown here because prophage detection is delegated to geNomad and evaluated separately. **(B)** Cross-genus generalization. Each of 13 experimentally validated tailocin producers, spanning five genera, was re-scored with its entire genus held out from training. Filled squares mark detection by PhageTAILor (genus held out) and by TattleTail. The tailocin subtypes are shown in parentheses. R denotes “rigid” R-type tailocins, F denotes “flexible” F-type tailocins, and R+F denotes tailocin regions that have both R-type and F-type components. PBSX is a defective prophage region with some capsid/head component and DNA packaging gene without full phage function but has tailocin function due to the presence of genes essential for tailocin function. PhageTAILor recovers 12/13 producers (0.92), missing only *Bacillus subtilis* (PBSX), whereas TattleTail recovers 2/13 (0.15), specialized on the two *Pseudomonas aeruginosa* strains its pyocin model was built for. **(C)** Head-to-head tailocin detection on the held-out curated benchmark (31 literature-confirmed producers and 5 negative controls). PhageTAILor (tailocin binary head at 0.5) is compared with TattleTail and geNomad-standalone (virus_score ≥ 0.5); bars and printed values give precision, recall, and F1. **(D)** Per-strain agreement between predicted and expected class on the labeled benchmark strains, stratified by the nearest-other-ANI confidence band (high ≥ 95, medium 90–95, low < 90, no match). n above each bar is the number of strains per band.

**Table 1.**
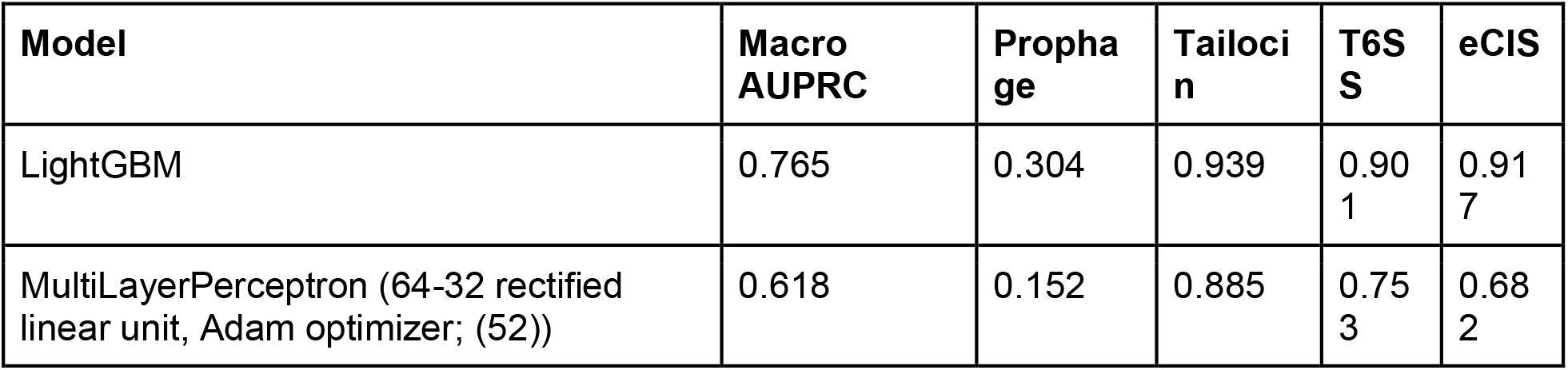
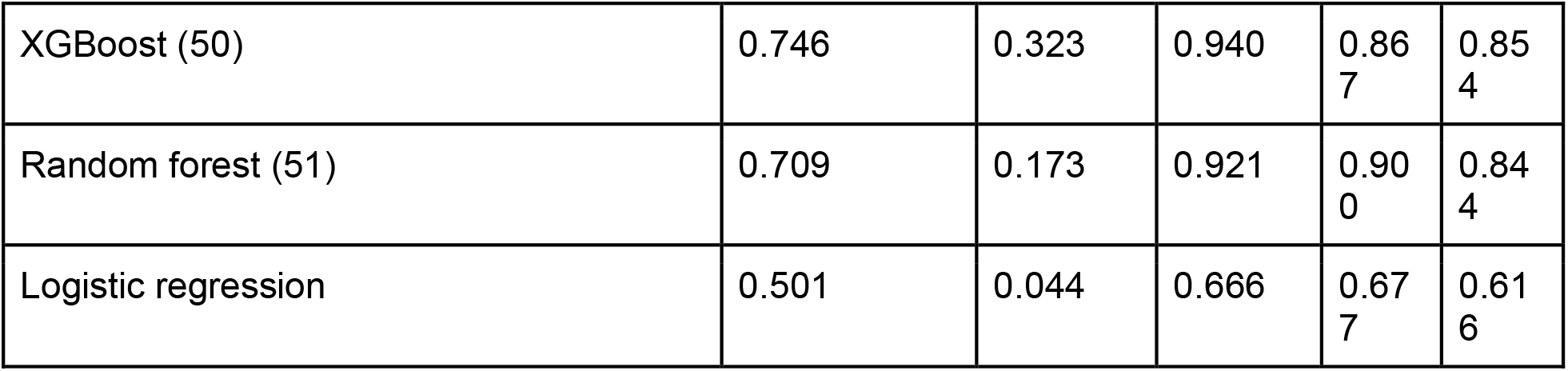
Model comparison on the same feature matrix. Per-class AUPRC (one-vs-rest), computed from out-of-fold predictions under genome-grouped 5-fold cross-validation on the identical 74-feature matrix. Macro (prophage/PTE) is the mean over the prophage and 3 PTE classes; models are ordered by it.

#### Tier 2. Cross-genus generalization (leave-one-genus-out)

The mean leave-one-genus-out multiclass macro F1 was 0.577. The drop from the in-distribution per-region macro F1 of 0.775 to 0.577 measures the genuine cross genus generalization gap, which is what the ANI based confidence band flags at inference. Cross-genus generalization for tailocins was evaluated directly in Tier 4.

#### Tier 3. Held-out cross-clade benchmark

On the held-out cross-clade benchmark, the tailocin binary head, max-pooled per genome and thresholded at 0.5, classified the 31 expected-positive strains together with 5 negative controls at precision 0.97, recall 1.00, F1 0.98, and AUPRC 0.97. Because ∼86% of this operating-point panel are true tailocin producers, its AUPRC is inflated—even a classifier that predicts every strain as positive would achieve a high score. To obtain a composition-independent estimate of precision, we instead evaluated the model on the clean-negative panel in Tier 4. Evaluated using the multiclass head, eCIS prediction achieved a precision of 0.83, recall of 0.62, and F1 of 0.71, with an AUPRC of 0.77 across eight positives, including PVC-type eCIS. Because both the eCIS labels and the eCIStem detector rely on the same reference catalog, these metrics measure the recovery of reference-like eCIS rather than true cross-clade generalization. Expanding the cross-clade eCIS positive set remains a priority for future work.

#### Tier 4. Clean-negative precision panel and cross-genus recall

On the clean-negative panel, whose composition is given in Supplementary Table S1, the deployed tailocin head reached precision 0.81 at recall 1.00, with no false positives on the hard same clade negatives or the structural negatives. The only false positives were three prophage bearing genomes, where prophage tail genes resemble tailocin structural genes. In the leave-one-genus-out recall evaluation—where each producer was scored with its entire genus excluded from training—PhageTAILor recovered 12 of 13 experimentally validated tailocins. This included all seven non-aeruginosa *Pseudomonas* strains and three of four remaining genera. By contrast, TattleTail detected only 2 of 13, identifying solely the *Pseudomonas aeruginosa* strains because its pyocin model does not generalize outside that species (Figure 2B; (2)).

### Algorithm Comparison

To confirm that predictive performance stems from the feature matrix rather than a specific algorithm, we benchmarked five classifier families—LightGBM, XGBoost, Random Forest, a multilayer perceptron, and logistic regression—on the identical 74-feature matrix using genome-grouped cross-validation (Figure S2). The two gradient-boosted decision tree architectures performed best. LightGBM had the highest macro AUPRC across the PTE and prophage classes at 0.765, followed by XGBoost at 0.746, Random Forest at 0.709, the MLP at 0.618, and logistic regression at 0.501. LightGBM and XGBoost were effectively tied on tailocin at 0.939 and 0.940, LightGBM led on T6SS at 0.901 versus 0.867 and on eCIS at 0.917 versus 0.854, and XGBoost was marginally better on the hardest class, prophage, at 0.323 versus 0.304. The close margin between the two boosted tree methods, set against the large gap to the others, indicates that the gradient boosted tree family rather than any single implementation drives performance on this feature matrix. We selected LightGBM due to its marginal performance advantage, faster training times, and native support for categorical features.

### Comparison with Other PTE-Prediction Tools

To determine whether a *Pseudomonas*-specific pyocin model or a broader viral detection tool could match our pipeline’s cross-clade generalization, we benchmarked PhageTAILor against TattleTail (2) and geNomad (5) on a literature-curated set of 41 strains, including 33 tailocin-positive isolates (Table 2). PhageTAILor attained the highest F1 of the tools compared at 0.955. Its recall of 0.970 matched geNomad standalone and exceeded TattleTail at 0.879, while its precision of 0.941 sat between TattleTail, which reached 1.000 at the cost of the lowest recall, and geNomad at 0.865. PhageTAILor therefore matched geNomad’s sensitivity without geNomad’s false positive rate and improved on TattleTail’s *Pseudomonas*-restricted recall. The single false negative was the cross-clade *Aeromonas dhakensis* SSU strain, which yielded a raw tailocin head score near zero. Although the HMM tail detector and genome-level HMM modules generated strong signal—detecting 11 distinct tail cassette profiles across 16 genes—these hits were fragmented across more than 70 draft contigs. Consequently, no single scaffold met the minimum threshold of ≥ 3 hits spanning ≥ 2 distinct profiles, and the per-genome HMM signal was insufficient to cross the decision boundary. The two false positives were prophage-bearing negative controls (*Escherichia coli* K-12 MG1655 and *Thermus thermophilus* HB8). In these strains, prophage structural genes mimicked tailocin tail modules, illustrating the same prophage-driven false-positive mode quantified on the Tier 4 clean-negative panel. In-distribution T6SS performance is detailed above, while prophage candidates are generated internally via geNomad rather than evaluated as a benchmark target.

**Table 2.**
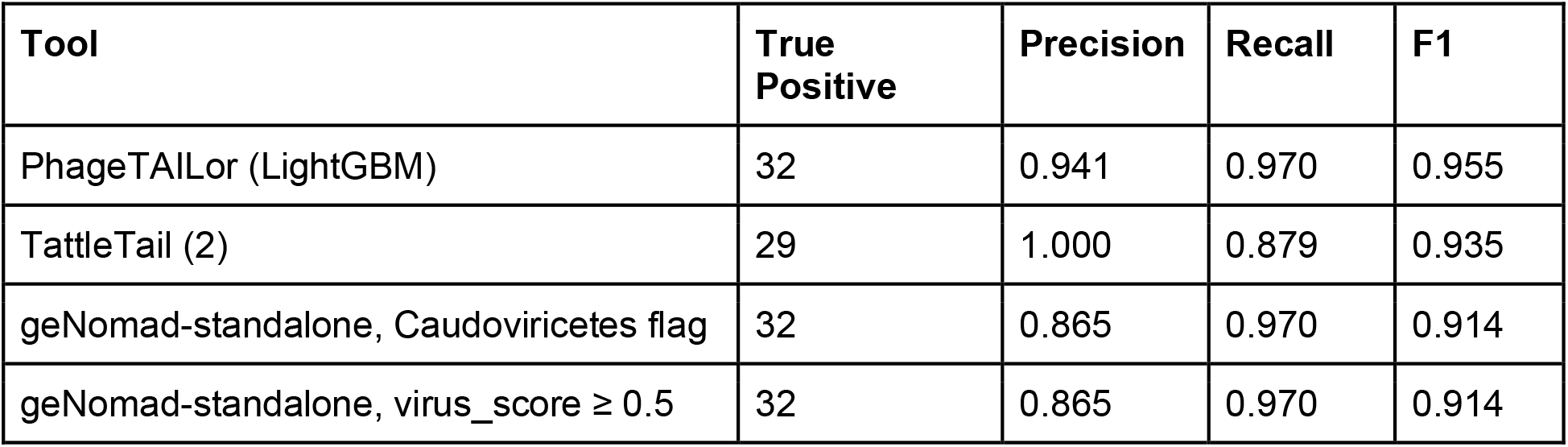
Head-to-head tailocin classification on the 76-strain held-out benchmark (33 positives). Metrics are computed on the 41 literature-labeled strains — 33 tailocin producers and 8 non-producers (2 eCIS, 5 negatives, 1 putative). PhageTAILor is scored with the deployed phylo-free model at its production operating point: the raw tailocin_binary head, max-pooled per-genome, thresholded at 0.5 (the isotonic binary-head calibrators are disabled in deployment, so the raw head is the served score). TattleTail and geNomad use each tool’s default operating point.

### Case study: PTE and prophage distribution across root-, leaf-, plant-, and soil-associated bacteria

We deployed PhageTAILor across 7,925 quality-filtered bacterial isolate genomes, partitioned by isolation site metadata into soil (n = 4,018), root-associated (n = 2,511), general plant-associated (n = 846), and leaf-associated (n = 550) niches. Running all five sequence and HMM detectors de novo at this scale fully populated every per-region feature block, enabling an unconstrained detection sweep with all PTE classes and prophages predicted directly by their deployed heads. Across the cohort, T6SS was the most prevalent class (43% of genomes), followed by tailocins (26%), prophages (21%), and eCIS (6%), with each class enriched within distinct GTDB orders (Figure 4).

Unadjusted detection rates varied markedly across niches (Figure 3C). Tailocin detection rose from 21.0% in Soil to 31.1% across the pooled plant niches, with a crude odds ratio of 1.69 at p = 2 × 10⁻²⁴, most strongly in bulk plant-associated bacteria at 41.5% and intermediately in the root-associated and leaf-associated compartments at 27.4% and 31.6%. T6SS followed the same pattern, rising from 37.9% in Soil to 48.8% in the pooled plant niches, with a crude odds ratio of 1.56 at p = 2 × 10⁻²². Prophage was weakly elevated at 22.3 versus 19.7%, with a crude odds ratio of 1.17 at p = 5 × 10⁻³, and eCIS showed the opposite soil-biased distribution at 8.4% in Soil versus 4.2% in the pooled plant niches, with a crude odds ratio of 0.48 at p = 2 × 10⁻¹⁴.

**Figure 3.**
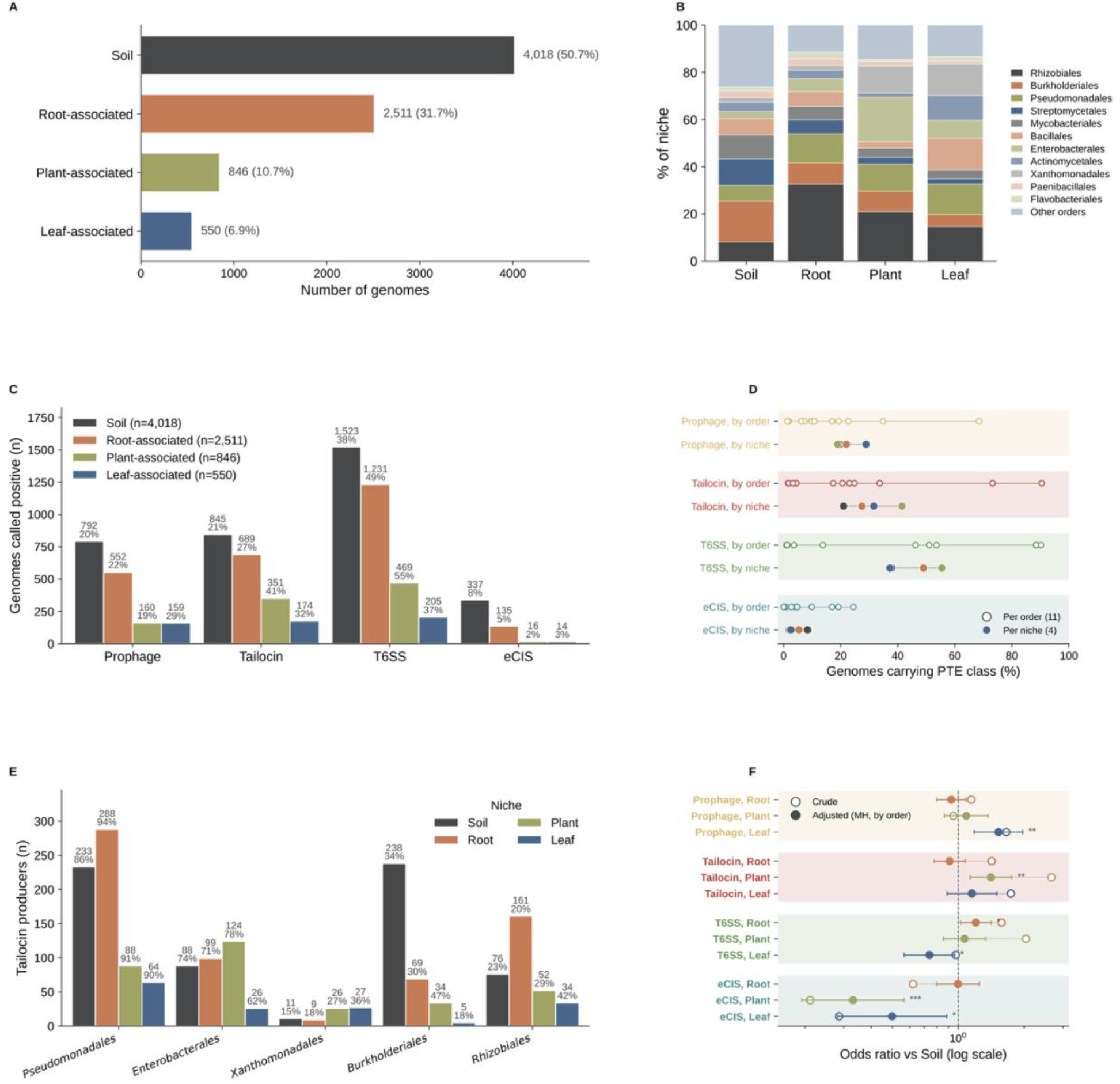
PTE and prophage distribution across 7,925 root-, leaf-, plant- and soil-associated bacteria, and their taxonomic composition. **(A)** Number of genomes per niche (Soil, n = 4,018; root-associated, n = 2,511; plant-associated, n = 846; leaf-associated, n = 550). **(B)** Per-niche taxonomic composition (stacked fraction of each niche by GTDB order); segment colors distinguish orders only (they do not encode niche). **(C)** Number of producer genomes per prophage or PTE class, by niche (bars are counts; each labeled with count and within-niche percentage), using the raw binary-head score ≥ 0.5 for tailocin and T6SS and multiclass probability ≥ 0.5 for prophage and eCIS. **(D)** Per-niche and per-class crude odds ratios versus Soil (Fisher’s exact test, log scale; each point is one plant niche; dashed line at OR = 1; * p < 0.05, ** p < 10⁻⁵, *** p < 10⁻¹⁰, ns not significant). **(E)** Number of tailocin-producer genomes within order, by niche, for the 5 most tailocin-rich orders (bars are counts, labeled with count and within-(order × niche) percentage); conditioning on order shows carriage is near-saturated and flat within tailocin-rich lineages. **(F)** Crude versus taxonomy-adjusted odds ratios for pooled plant versus Soil, per class (adjusted = Mantel–Haenszel estimator stratified by GTDB order, Robins–Breslow– Greenland 95% confidence intervals). Every crude contrast collapse to OR ≈ 1 after adjustment, showing the niche signal is almost entirely compositional.

However, these raw differences were largely driven by taxonomic composition. Plant-associated niches were heavily enriched for a few Pseudomonadota orders that harbor the majority of tailocins and T6SS (Figures 3B and 4). Indeed, PTE and prophage prevalence varied far more across GTDB orders than across ecological niches (Figure 3D). Therefore, we re-estimated each niche contrast as a Mantel Haenszel odds ratio stratified by GTDB order (Figure 3F). The broad plant-versus-soil differences disappeared across all classes (adjusted odds ratios = 1.0–1.1; p > 0.1), confirming that the apparent plant enrichment was entirely driven by taxonomic composition. However, several specific niche associations remained significant even after adjustment. Tailocins remained enriched in bulk plant-associated bacteria with an adjusted odds ratio of 1.41 at p = 3 × 10⁻³, prophages in the phyllosphere with an adjusted odds ratio of 1.53 at p = 2 × 10⁻³, and eCIS remained depleted in plant-associated bacteria with an adjusted odds ratio of 0.33 at p = 3 × 10⁻⁵. All three associations remained significant after Bonferroni correction across the twelve niche-specific tests. Weaker, nominally significant effects included T6SS enrichment in the rhizosphere at an adjusted odds ratio of 1.20 and reduced T6SS and eCIS in the phyllosphere at 0.74 and 0.50. The pooled analysis appeared null because it averaged heterogeneous—and occasionally opposing—niche-specific effects; for instance, T6SS was enriched in the rhizosphere but depleted in the phyllosphere. Conditioning on order confirmed this taxonomic confounding. Within tailocin-rich orders like Pseudomonadales, carriage was nearly saturated and flat across habitats (Figure 3E), while per-order phylogenies demonstrate that PTE carriage tracks lineage rather than environment (Figures 6 and S12). Thus, while crude niche associations were largely artifacts of taxonomic sampling, several biologically sensible, taxonomy-independent niche effects persisted. This highlights that our genome-scale predictions are sensitive enough to both unmask taxonomic confounding and isolate genuine biological signal.

**Figure 4.**
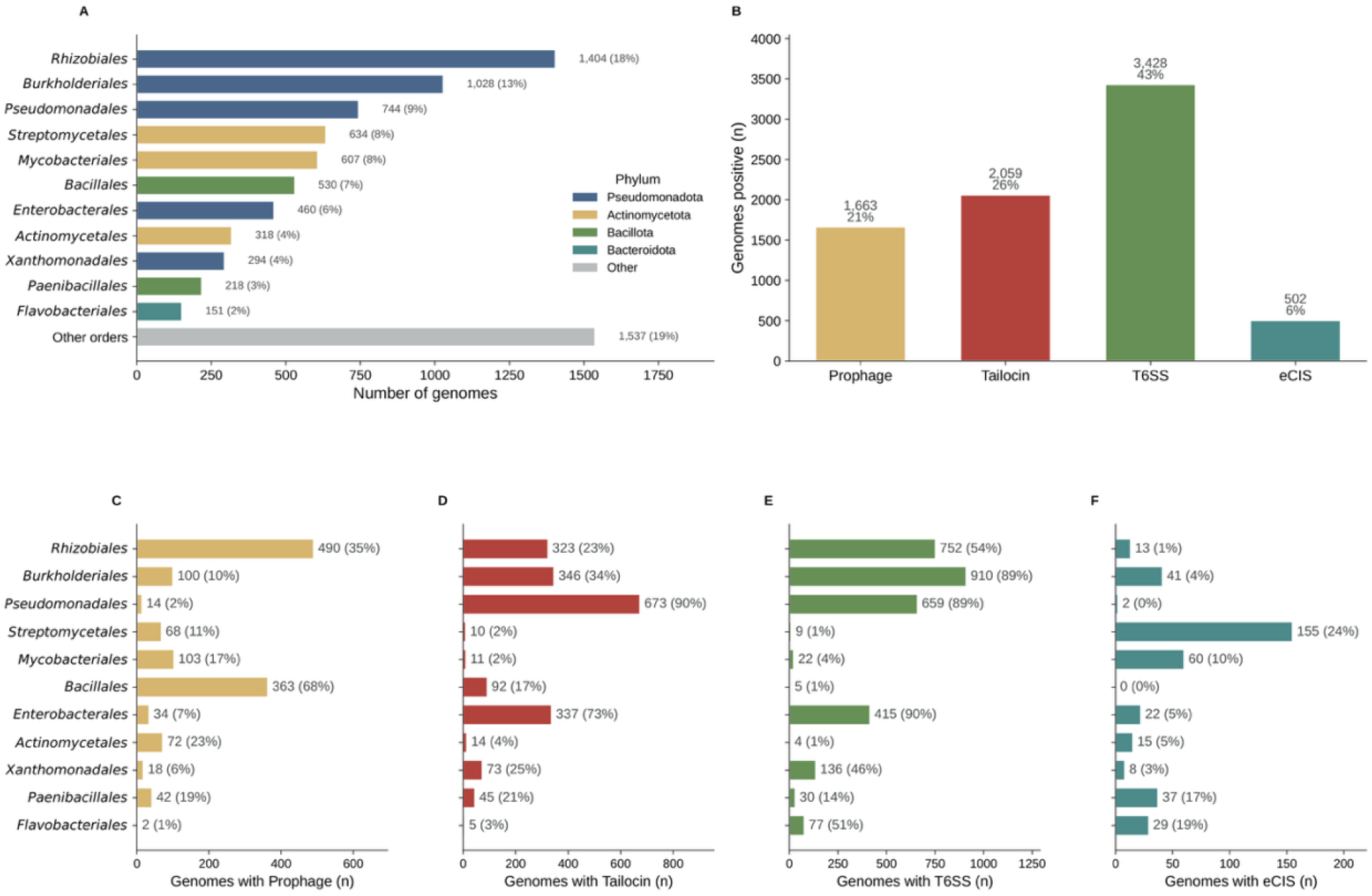
Taxonomic landscape of prophage or PTE classes across the 7,925-genome plant- and soil-associated cohort. **(A)** Most abundant GTDB orders across the 7,925 genomes, colored by phylum (remaining orders pooled as “Other orders”). **(B)** Total number of genomes carrying each prophage or PTE class across the full 7,925 genome set (each bar labeled with count and percentage of the 7,925-genome set). **(C–F)** Number of genomes carrying each prophage or PTE class within each GTDB order from the same deployed calls; each bar is labeled with count and within-order percentage, orders share one top-to-bottom ordering (by total genome count) across the four panels, and bars are colored by prophage or PTE class. **(C)** Prophage. **(D)** Tailocin. **(E)** T6SS. **(F)** eCIS.

**Figure 5.**
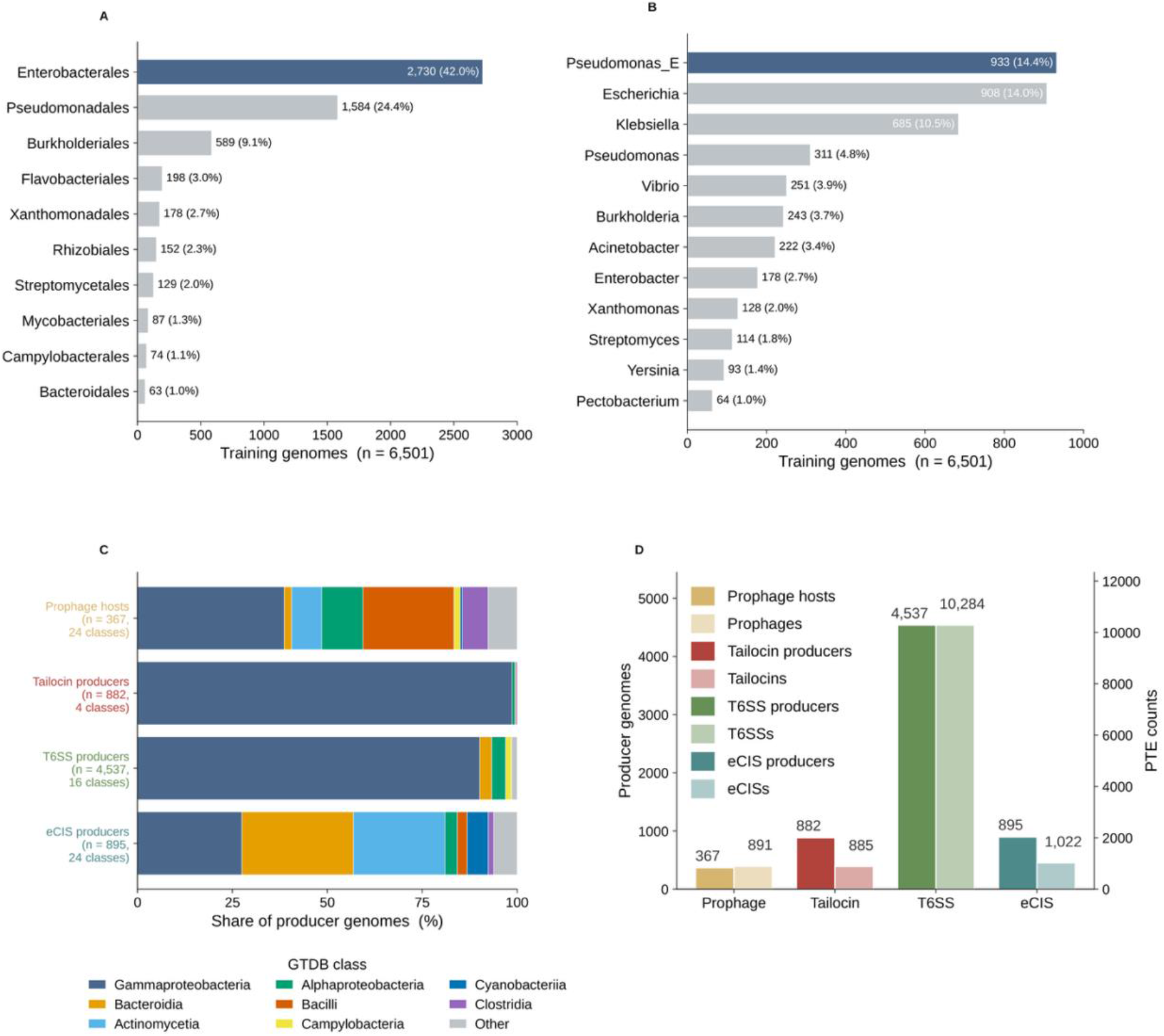
Taxonomic affiliation and prophage or PTE numbers of training genomes. **(A)** Top 10 most abundant orders. **(B)** Top 12 most abundant genera of training genomes. **(C)** Class composition of training genomes, grouped by prophage or PTE types. **(D)** Number of producer genomes and PTEs or phages of each prophage or PTE class in training genome data.

**Figure 6.**
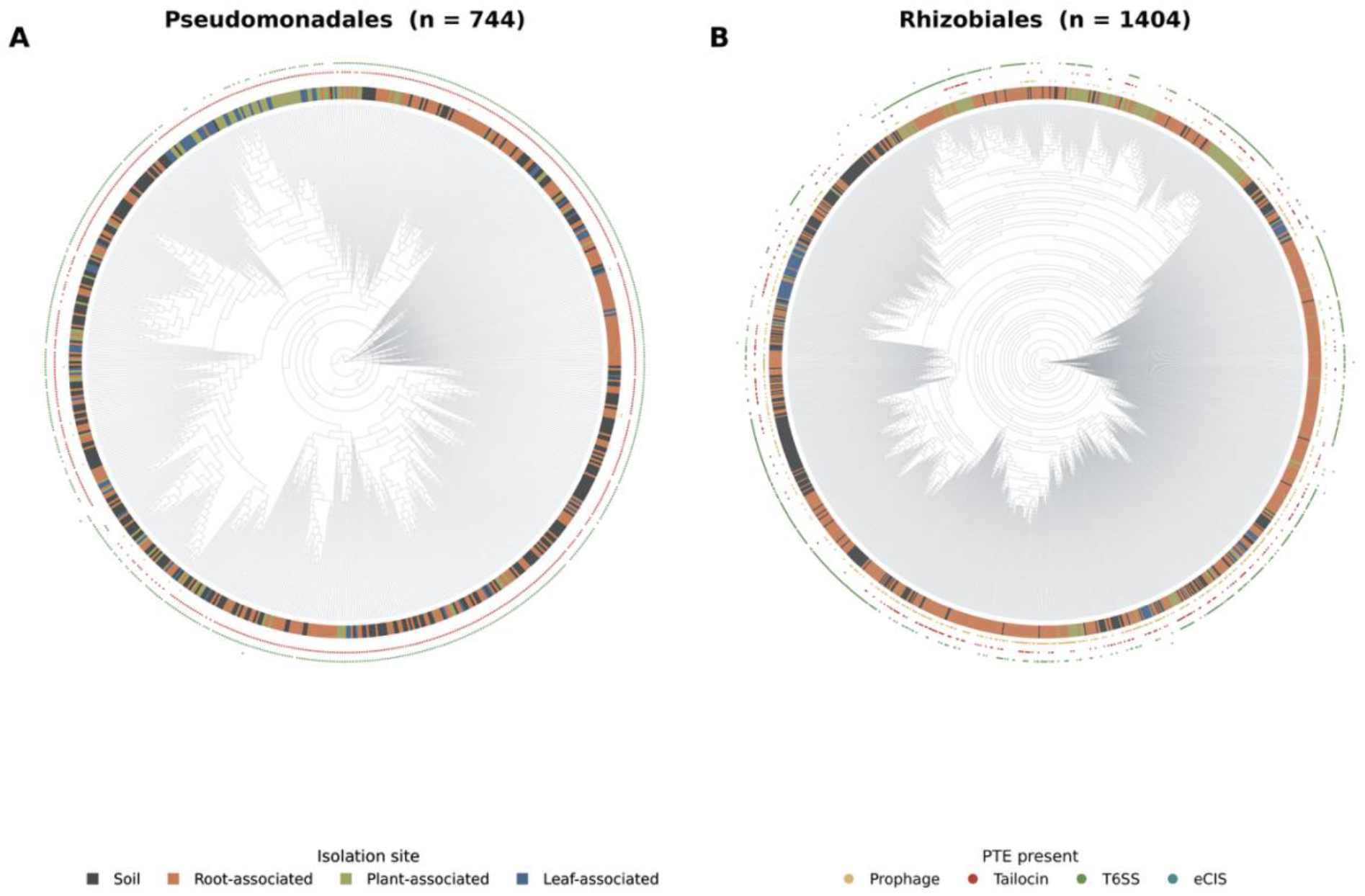
Prophage or PTE occurrence is governed by phylogeny, not isolation niche. Circular cladograms (aligned tips) of two contrasting GTDB orders; the inner ring colors each genome by isolation niche and the four outer dot rings mark presence of each prophage or PTE class (deployed-model calls). Per-tip names are omitted for legibility (labeled per-order versions in Fig. S12). **(A)** Pseudomonadales (n = 744): tailocin and T6SS are present in nearly every genome across the entire tree while the niche ring is fully intermixed — prophage or PTE carriage is a near-universal lineage trait, independent of niche. **(B)** Hyphomicrobiales (n = 1,404): Prophage or PTE carriage is instead clade-restricted, confined to sub-clades, again with niche scattered across the tree. In both, prophage or PTE presence tracks phylogenetic position rather than niche — consistent with the collapse of the niche signal after taxonomy adjustment (Fig. 3F).

## DISCUSSION

PhageTAILor addresses key limitations of single-element PTE and prophage detection tools, as well as the constraints of currently available tailocin prediction methods. PhageTAILor detects and classifies prophages along with three contractile PTE classes—tailocins, T6SSs, and eCISs—within a unified, genome-scale pipeline. Crucially, it retains raw evidence from each detector as a primary output, allowing users to independently inspect and reuse detection and classification data. Furthermore, it provides an automatic pipeline to detect tailocins across various taxa of bacteria, filling in the gaps left by tailocin prediction approaches that are either not fully automated or specializing in the most widely studied tailocin producer taxa.

PTE and prophage detection of PhageTAILor allows generalization beyond the training dataset, highlighting the practical value of this pipeline as a PTE and prophage predictor by working well on lineages outside of the curated reference sets that are heavily Pseudomonadota-biased. On the held-out 76-strain benchmark, PhageTAILor attains the highest F1 (0.955) of the tools compared, matching geNomad’s recall without its false-positive rate and exceeding TattleTail’s *Pseudomonas-*specific recall. In the more stringent leave-one-genus-out evaluation, PhageTAILor recovered 12 of 13 experimentally validated tailocins, compared to just 2 of 13 for TattleTail. This advantage is largely driven by our HMM tail-profile library, which provides sequence-divergence-tolerant detection where MMseqs2 hits saturate on highly diverged hosts.

To prevent data contamination, we deliberately designed PhageTAILor to operate without phylogenetic features, enforcing an explicit leakage budget. We excluded taxonomy and phylogenetic placement features not because they reduce in-distribution performance—they increase macro F1 by ∼0.086—but because they cannot be reliably reproduced at inference time and primarily represent genus memorization rather than transferable biological signal. Our leakage decomposition quantifies this trade-off explicitly: macro F1 declines from 0.886 to 0.861 when transitioning from random to genome-grouped folds, and further to 0.775 when removing phylogenetic features entirely. The remaining cross-genus generalization gap (0.775 to 0.577 under leave-one-genus-out cross-validation) is directly flagged to users by our inference-time ANI confidence score. Ultimately, we argue that reliable in-distribution predictions, transparent reporting of generalization limits, and instance-level confidence bounds are far more useful in practice than inflated headline metrics that rely on data leakage. This modeling challenge stems directly from the extreme lineage skew of PTE and prophage distributions across bacterial clades (Figures 4C–F and S11). Each group of PTEs and prophages tend to be found specifically more widespread in less than a handful of classes, partially due to the genuine uneven occurrences of them across bacterial taxa, and partially due to the shortage of studies in specific PTEs and prophages in certain lineages. For example, T6SSs gene clusters are classified as type i, ii, and iii, where type i are mostly carried by the Pseudomonadota phylum, type ii found in the *Francisella* pathogenicity islands, and type iii found in the Bacteroidota phylum (24). Considering that tailocins and eCIS are relatively newer, rapidly expanding areas of research, there may be additional lineages harboring these classes of PTEs that are yet to be reported.

Selecting operating points on a per-class basis—sourcing tailocins and T6SS from dedicated binary heads while deriving eCIS and prophage calls from the multiclass head—allows a single model architecture to effectively handle four classes with vast differences in baseline prevalence.

Our case study using 7,925 root-, leaf-, plant-, and soil-associated bacteria shows that lineage may play more important roles than niche in PTE and prophage carriage, reproducing the niche-enrichment patterns that are widely found in other comparative-genomics surveys (Figure 3D and Figure 6).

Because these three correlational trends remain significant after adjusting for taxonomy, they represent within-lineage associations rather than compositional artifacts. The phyllosphere is a harsh, dynamic environment characterized by intense UV radiation, desiccation, and frequent thermal and nutrient fluctuations (27). These environmental stressors are classic triggers of the bacterial SOS response and prophage induction; consequently, high phage pressure and active phage–host dynamics on leaf surfaces favor lysogeny (28, 29). The temperate life cycle is advantageous when host populations are spatially patchy and stress or phage encounters are frequent, conferring superinfection immunity to host cells (29, 30). Also, prophages frequently carry morons, which are extra genes that provide fitness advantage, that include stress-tolerance genes and often the toxins used in competition (31). These may be advantageous in the competitive environment on the exposed surface of the leaf. Either way, more prophages in one of the most stress-exposed plant compartments is plausible.

Selection for interbacterial competition during host colonization may explain the importance of interbacterial competence in a plant-associated environment with high bacterial density, consistent with tailocins being well documented in plant-associated *Pseudomonas* as colonization and competition tools. The eukaryotic predators or hosts that eCIS target are abundant in soil as well as in plant-associated environments. Within plant-associated lineages of bacteria, the strains that keep these costly machineries are likely those still engaged in soil and eukaryote interactions, and strains specialized to plant environment tend to lack eCIS.

Prophages and tailocins mediate interbacterial competition and stress tolerance well-suited to the densely populated, stressful plant surface, whereas eCIS systems predominantly target eukaryotic hosts abundant in soil (32, 33). Our results indicate that taxonomy-adjusted niche preferences reflect the target organisms of each prophage or PTE class rather than general habitat properties. Importantly, these interpretations are derived from modest, taxonomy-adjusted presence/absence associations rather than expression or functional assays. Experimental validation—such as induction assays, competition experiments, and effector-target identification—will be necessary to confirm these functional roles. Also, methodological limitations are present in our work. Unadjusted carriage-rate comparisons across niches are easily confounded by the taxonomic composition of sampled genomes, requiring a tool sensitive enough to separate raw ecological signals from underlying lineage bias.

Our pipeline runs in about 3 minutes per 5-Mb genome on 8 cores and is fully modular (any single detector can be run on its own). Since PhageTAILor reuses precomputed detector outputs when present, it scales from single genomes to the thousands-of-genomes surveys as we have exemplified using our case study.

The primary remaining limitations involve cross-clade generalization and reference dependence. The training dataset is heavily skewed toward Pseudomonadota (83.5% overall; 86% of tailocin producers are *Pseudomonas*), meaning detection sensitivity is highest on familiar lineages. Predictions on distant taxa (e.g., Cyanobacteria, Spirochaetota) should therefore be interpreted alongside the nearest_other_ani confidence metric. Consequently, the cross-clade benchmark remains sparse for certain classes; with only eight held-out cross-clade eCIS positives, per-class eCIS precision and recall cannot be reliably estimated at this evaluation tier. While our HMM library enables the detection of divergent tail loci, remaining failures on cross-clade tailocins stem from spatial organization rather than sequence divergence. For example, on the sole benchmark false negative (*Aeromonas dhakensis* SSU), robust HMM signal (11 tail-cassette profiles across 16 genes) was fragmented across 70 draft contigs, preventing any single scaffold from meeting the detection threshold. Two fundamental limitations remain. For three classes, the positive labels and their corresponding detectors derive from the same reference source, introducing a potential circularity concern. Leave-one-genome-out and leave-one-genus-out controls confirm that detection is not self-dependent—every eCIS genome is successfully recovered even when its own operons, or those of its entire genus, are removed. Nevertheless, a conceptual boundary remains: the model identifies elements matching known reference families, meaning genuinely novel families lie beyond its scope and remain undertested due to the small eCIS reference set. Furthermore, because the binary classification heads integrate features across multiple detectors, passing a zero-filled rather than fully computed feature block can cause over-prediction (e.g., miscalling tailocins across most candidate regions). While running all detectors de novo circumvents this issue, individual heads are not yet robust to missing input blocks. Finally, element detection remains inherently easier than target identification; tail fibers govern host specificity and diversify rapidly, making host-range prediction significantly more challenging than locus detection.

These constraints define our primary development priorities. The most pressing benchmark objective is expanding the cross-clade eCIS positive dataset to 10–20 strains to stabilize held-out metrics, alongside incorporating diverse cross-clade training positives to extend coverage across underrepresented taxa. To recover tailocin loci split across fragmented assemblies, a fragmentation-aware aggregation step could pool HMM hits across neighboring contigs prior to region calling. Complementing this with structural search algorithms—such as Foldseek queried against AlphaFold or ESM models of canonical tail proteins (34–36) —would anchor highly diverged fibers on broken scaffolds, providing a logical bridge from tail fiber detection to host-range identification. On the model architecture side, restricting each binary head exclusively to its corresponding detector’s features (feature masking) would ensure robustness when users execute only a subset of detectors.

## MATERIALS AND METHODS

### Pipeline Overview

PhageTAILor predicts prophages and three classes of phage tail like elements, namely tailocins, T6SS, and eCIS, in bacterial genomes through a Snakemake workflow (37). The codebase comprises a model construction workflow (Part A), run once per model release, and a user inference workflow (Part B), executed by end users on their own genomes. Both branches share a unified gene caller, reference database set, 74-feature per-region vector, and candidate union across six detectors. Standardizing these components ensures that user genome predictions remain directly comparable to the training corpus.

### Software, versions, and parameters

All software tools, pinned versions, and non-default parameters used by PhageTAILor are listed in Table 3; complete Conda environment specifications are provided in the repository. In brief, genes are called with pyrodigal-gv v0.3.2 backed by Pyrodigal 3.6.3 using ‘-p meta’ (5, 38, 39). Sequence searches were conducted using MMseqs2 (v14) (40) with search -e 1e-5 -s 5.7 --max-seqs 5 against the SecReT6, eCIStem, and tail gene databases, and -s 5 --max-seqs 3 against PHROGS. Profile HMM searches were run using HMMER (v3.3) (41) with hmmsearch -E 1e-5 against our custom tail HMM library. Proviruses were identified using geNomad (v1.7.4) (5) with end-to-end --relaxed. Nearest-neighbor training ANI was computed using skani (v0.3.1) (42) with search --min-af 50. Taxonomic assignments were generated using GTDB-Tk (v2.6.1) against GTDB release r226 (43), and genome quality was restricted to high-quality drafts using CheckM2 (v1.0.1) (44) (completeness ≥ 90%, contamination ≤ 5%). The classifier was trained using LightGBM (v3.3) (26) (num_iterations=500, learning_rate=0.1, num_leaves=64, max_depth=6, min_data_in_leaf=20, bagging_fraction=0.8, feature_fraction=0.8, data_random_seed=1). The entire pipeline was orchestrated using Snakemake with rule-specific Conda environments (37).

**Table 3.**
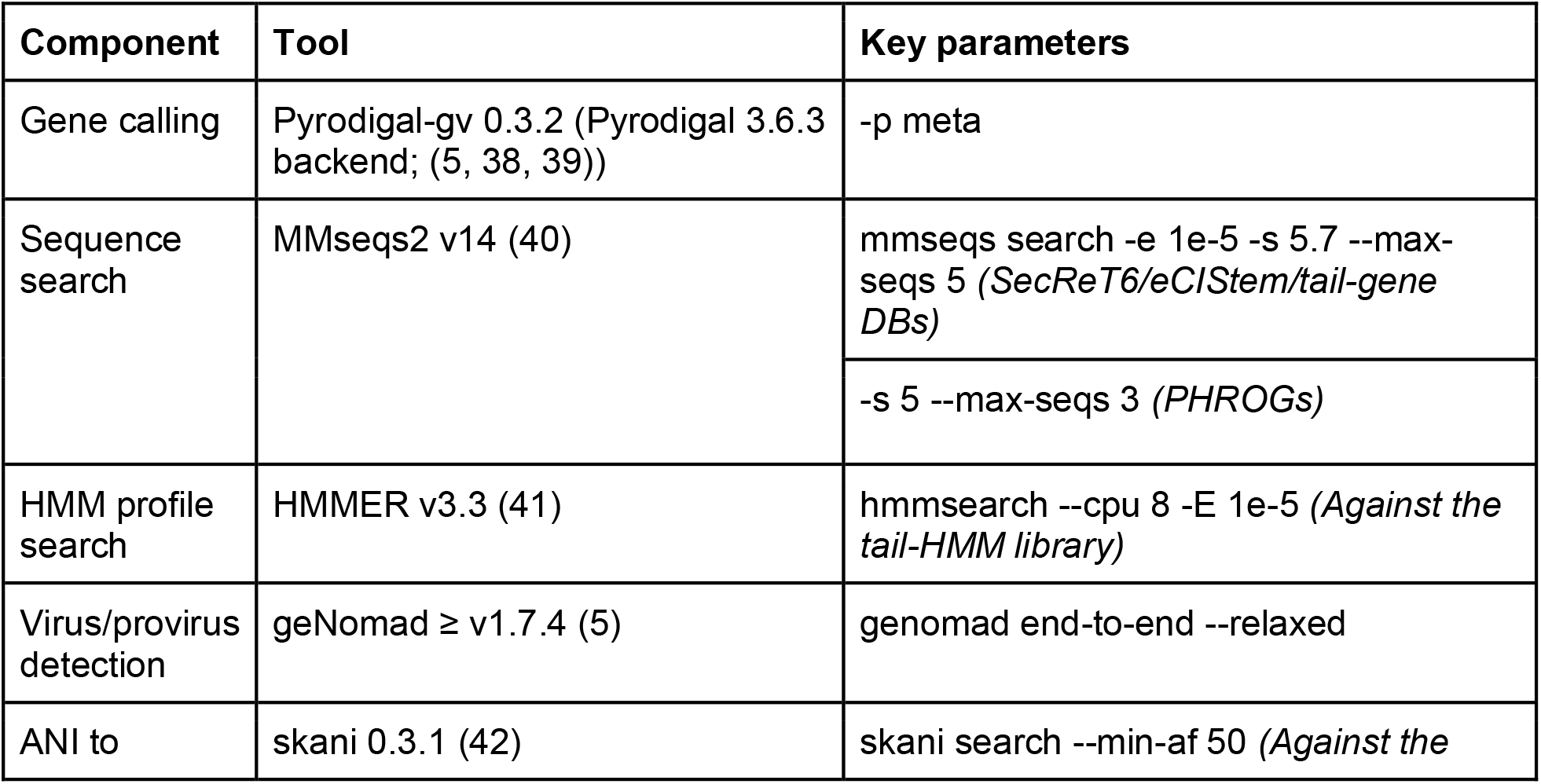
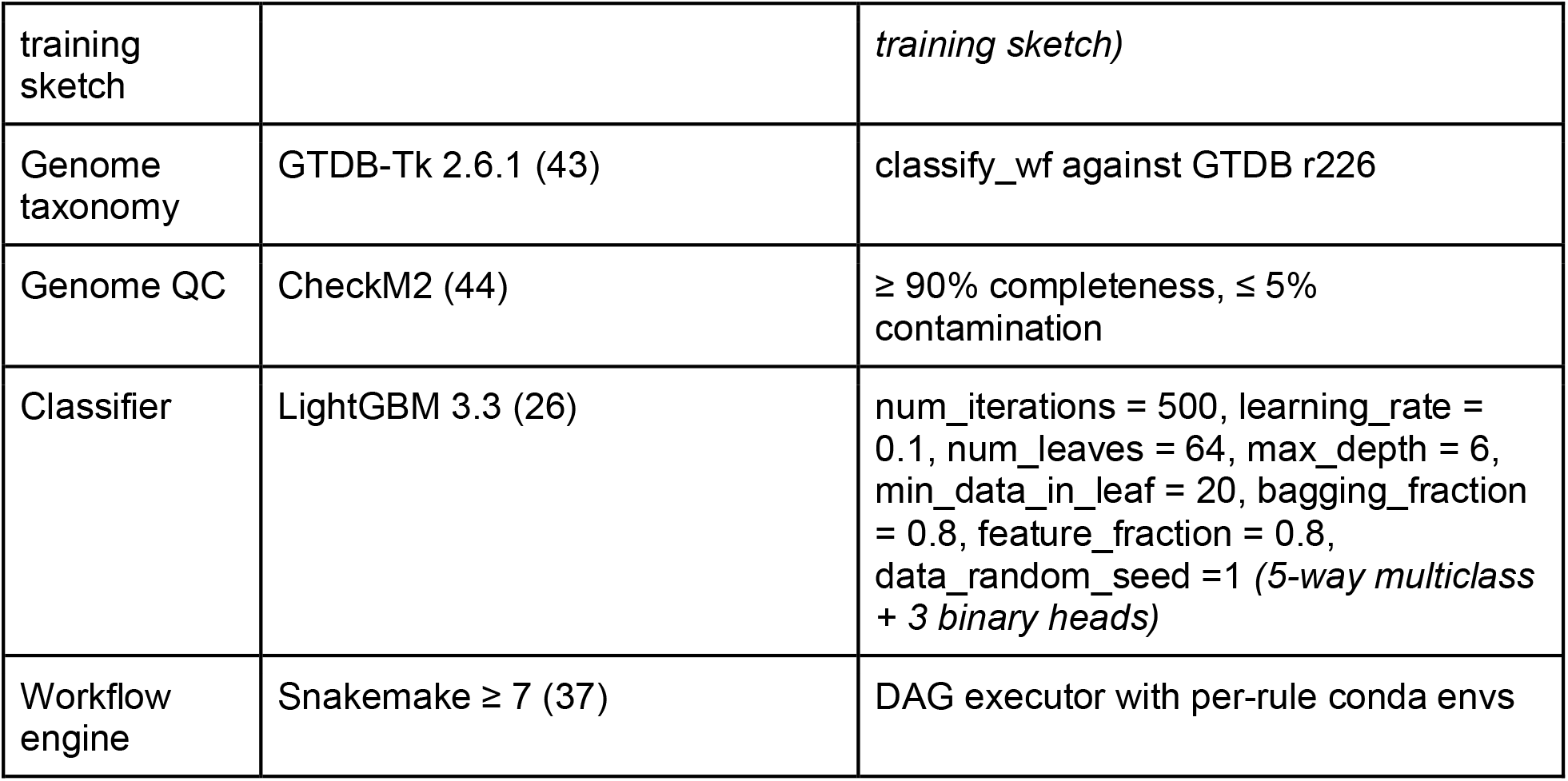
Software used in PhageTAILor.

**Table 4.**
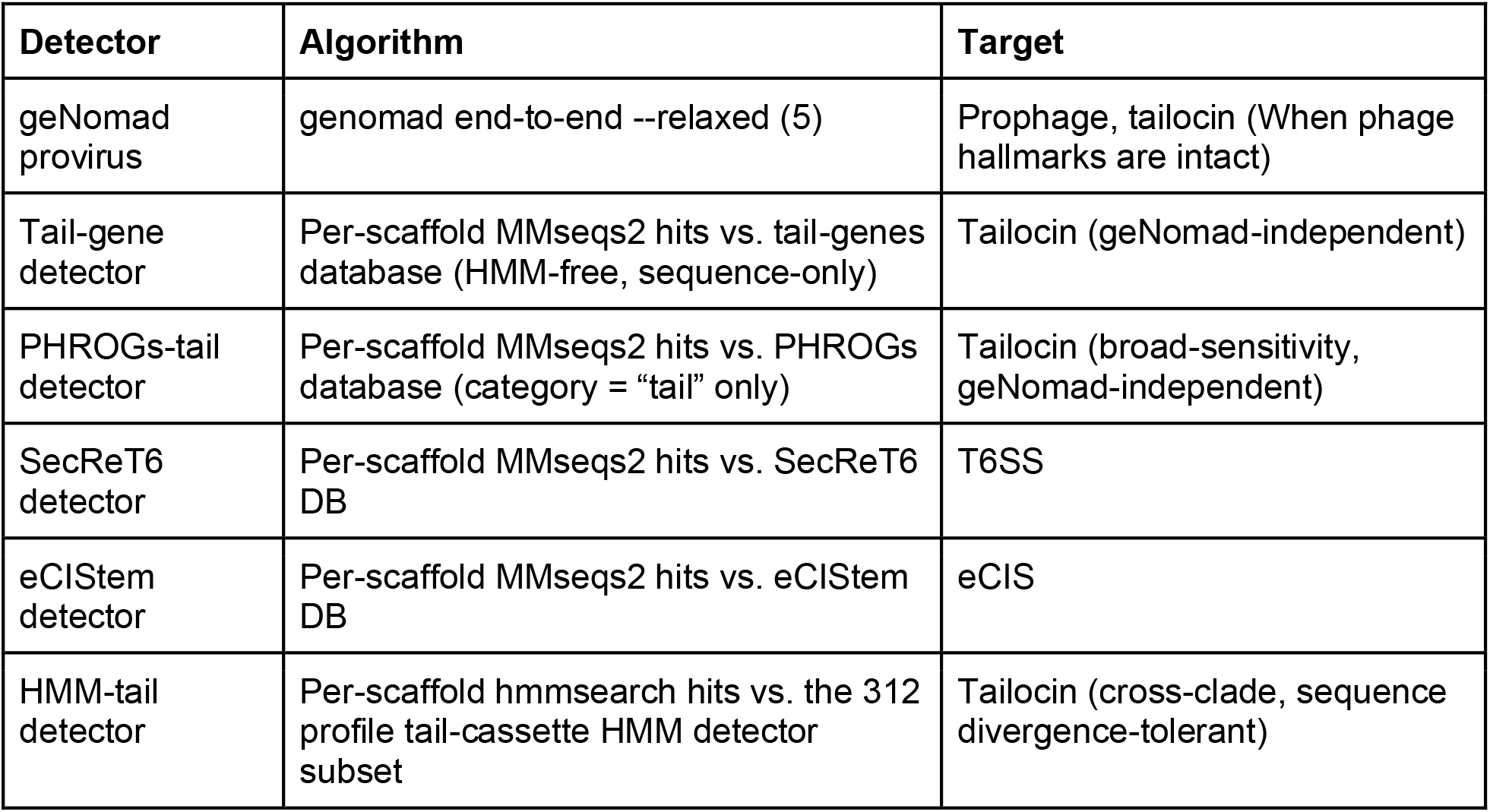
The 6-detector candidate region union.

### Part A. Model construction

#### Training-genome curation

We assembled a training corpus of 6,501 bacterial genomes spanning prophages and 3 PTE classes. Annotation evidence for each genome is represented as a curated coordinate interval defined by scaffold, start position, end position, and element type. In total, the coordinate dataset comprises 13,082 intervals: 10,284 T6SS, 1,022 eCIS, 891 prophage (708 Phage1 and 183 Phage2), and 885 tailocin loci aggregated from NCBI RefSeq (45, 46) and JGI IMG databases (47). Tailocin coordinates were manually curated from primary literature describing experimentally validated loci, with source publication DOIs recorded for each entry. Prophage coordinates were derived from the curated dataset of Mageeney et al. (48), with Phage1 and Phage2 tiers merged into a single prophage class during model training. T6SS coordinates were sourced from SecReT6 (24)., and eCIS coordinates were obtained from eCIStem (3), re-anchored to their original target genome assemblies.

Every producer genome was required to have at most 200 scaffolds, ≥ 90% CheckM2 completeness (44), and ≤ 5% contamination, and was then dereplicated within each PTE producer group at 99.995% pairwise ANI with skani (42). Genome dereplication removed 726 genomes from the T6SS dataset and 16 from the prophage dataset (including one genome redundant in both); the tailocin and eCIS sets were already non-redundant, and no removed genome represented the sole instance of a tailocin or eCIS locus. Because dereplication was executed within each class, near-identical genomes were both retained if they harbored different PTE classes on a shared chromosomal background. Any residual strain similarity introduced by this approach is controlled during model evaluation: all cross-validation folds are partitioned by genome to ensure no assembly spans training and test sets, and primary generalization metrics rely on leave-one-genus-out splits, which explicitly separate closely related strains belonging to the same genus.

#### Taxonomic Classification and ANI calculation

Taxonomies for all training-set genomes were assigned using GTDB-Tk (v2.6.1) classify_wf against GTDB reference release r226 (43). The 6,501-genome training set spans 24 phyla, 42 classes, 102 orders, 214 families, 624 genera and 1,919 species. This training genome set comprised 5,431 Pseudomonadota, 416 Bacteroidota, 247 Actinomycetota, 119 Bacillota, 74 Campylobacterota, 54 Cyanobacteriota, 35 Bacillota_A, 29 Planctomycetota, 24 Desulfobacterota, 21 Myxococcota, 11 Desulfobacterota_F, and 13 other phyla with less than 10 genomes each (Figure 5). Additionally, a skani sketch of the 6,501 training genomes (4,164,667,441 bytes of total sketch database) is bundled with the model; this is queried at inference time to derive the nearest_other_ani confidence band.

#### Reference databases and homology search

Homology searches integrate five reference databases: SecReT6 (24), an eCIStem cluster set (3), a custom tail-gene sequence database, PHROGS profiles restricted to tail-related categories (9), and a 597-profile tail HMM library, alongside geNomad for provirus detection (5). A 312-profile subset corresponding to core structural modules (baseplate, sheath, tube, and related components) is dedicated to candidate-region nomination, whereas the complete 597-profile library generates both per-region and genome-level HMM features. The unified training protein set was concatenated with sample prefixed headers into a single MMseqs2 query database and searched against each sequence resource with the parameters in Table 3. Per query hits with a bitscore of ≥ 50 and ≥ 30% identity were retained and demultiplexed back to their source genome, and HMMER hits were retained at an E value of at most 1e-5.

#### Candidate region detection

Candidate regions are defined by taking the union of outputs from six individual detectors (five sequence-based and one HMM-based). Each detector aggregates colocalized hits along a scaffold into a discrete candidate region, a spatial grouping step that is distinct from the sequence clustering performed by MMseqs2 during initial homology searches. The four MMseqs2 colocalization detectors share one per scaffold gap merging rule. For each detector, candidate regions were nominated by retaining the top bitscore hit per query gene, sorting hits by start coordinate, and merging consecutive hits separated by an intergenic gap of at most 15 kb. A candidate region was emitted if the merged window contained ≥ 4 hits spanning ≥ 3 distinct components, clusters, or sequences. The PHROGS tail detector and the HMM tail detector applied a relaxed threshold requiring ≥ 3 hits across ≥ 2 distinct tail PHROGs or HMM profiles, respectively. These heuristic thresholds were adopted from the published SecReT6 workflow. Across all training genomes, the five sequence-based detectors emitted 549,926 candidate regions, including 372,584 from eCIStem, 111,029 from the tail gene set, 35,027 from geNomad, 16,076 from SecReT6, and 15,210 from PHROGS tail. The HMM tail detector populates feature blocks for these candidates during training and serves as an active sixth candidate source at inference, where it nominates HMM-only regions to recover divergent cross-clade tail loci.

#### Region labeling

Each candidate was labeled against the curated training intervals using a 50% reciprocal overlap rule, requiring both the intersection over the candidate length and the intersection over the truth length to be ≥ 0.5, implemented as an in-memory pandas routine. Conflicts between overlapping ground-truth intervals were resolved by maximizing the minimum reciprocal overlap, with unmatched candidates designated as hard negatives. This strict matching criterion maintains high positive-label purity, but penalizes prophages whose curated boundaries are inherently large and diffuse—providing additional rationale for delegating primary prophage detection to geNomad.

#### Per-region feature extraction

Each candidate region is encoded as a fixed 74-feature vector. This feature set captures region geometry and gene architecture (length, gene count, gene density, strand orientation consistency, and tail gene span); cluster-level signals across detectors (hit counts, distinct component counts, and top bitscores); category-specific PHROGS counts; geNomad output metrics (virus score, hallmark count, marker enrichment, and Caudoviricetes classification); and regional and genome-wide tail HMM module counts. To prevent phylogenetic shortcuts, the feature matrix explicitly excludes taxonomic assignment (label-encoded genus, family, and order), nearest-neighbor training ANI, and the number of close training genomes. The nearest-neighbor training ANI is calculated exclusively to establish inference-time confidence bands and is not supplied as an input feature to the model.

#### Classifier training

PhageTAILor emits four prediction heads trained with LightGBM 3.3 (26), a five way multiclass head over negative, tailocin, prophage, T6SS, and eCIS with softmax output, and three binary heads for tailocin, T6SS, and eCIS. All 4 heads share the 74-feature matrix. Class imbalance was addressed using positive class weighting equal to the negative-to-positive ratio for the binary heads, and inverse class-frequency sample weights for the multiclass head. We used raw output probabilities from the binary heads without post-hoc isotonic calibration, as calibration proved unstable under severe class imbalance and provided no performance gains at the deployed operating points. Hyperparameters, verified directly from the saved boosters, are detailed in Table 3.

#### Inference time confidence band

During inference, each genome is assigned an ANI-based confidence band derived from its maximum ANI to any training genome, computed using skani (42). Confidence tiers are categorized as high (≥95% ANI), medium (90% to 95% ANI), low (<90% ANI), or “no match” if no training genome meets the minimum alignment coverage. This confidence score is reported alongside every prediction and is not used as a model input feature.

#### Model evaluation

Per-class genome-level predictions apply a 0.5 probability threshold to the raw binary heads for tailocin and T6SS, and a 0.5 threshold to multiclass probabilities for eCIS and prophage, taking the maximum score across a genome’s candidate regions in each case. Model performance was evaluated across four tiers. Tier 1 is in-distribution genome-grouped five-fold cross-validation, in which folds were partitioned by genome using scikit-learn GroupKFold so no genome appeared in both training and test sets (49). Tier 2 is leave-one-genus-out cross-validation, which iteratively holds out every representative genome of a genus in turn. Tier 3 is an independent held-out cross-clade benchmark of 76 strains completely withheld from training, including 41 literature-annotated strains (33 of which are tailocin producers). Tier 4 is a clean-negative precision panel paired with a cross-genus recall test, detailed in Supplementary Table S1. This panel combines 18 curated tailocin-free genomes across three categories—experimentally confirmed non-producers, non-contractile bacteriocin producers lacking tailocins, and obligate intracellular or minimal genomes lacking contractile machinery—with 13 experimentally validated producers spanning five genera. Benchmark metrics are reported pooled across all confidence bands, and the case study is additionally stratified by band to confirm that raw niche contrasts are not an artifact of low-confidence predictions.

#### Algorithm comparison

To confirm that LightGBM is appropriate and that the model architecture is not a performance bottleneck, we evaluated four alternative classifiers: XGBoost (50), Random Forest (51), a multilayer perceptron (MLP; 64- and 32-unit hidden layers, ReLU activation, Adam optimizer) (52), and logistic regression, all implemented via scikit-learn (49). These models were trained on the identical 74-feature matrix using the exact genome-grouped folds as the production model. Classifiers were evaluated using per-class one-versus-rest area under the precision-recall curve (AUPRC) rather than F1 score, as threshold-based F1 metrics were unstable across folds under class-imbalance weighting, whereas AUPRC provides a threshold-free measure of regional ranking performance. Comparative results are detailed in Table 1.

#### Comparison with other PTE prediction tools

On the 76 strain held out benchmark, PhageTAILor was compared against published tailocin and prophage prediction tools, namely TattleTail (2) and geNomad standalone (5), using each tool’s default operating point. PhageTAILor was scored with the deployed phylo-free model at its production operating point, the raw tailocin binary head max-pooled per-genome and thresholded at 0.5. Metrics were computed on the 41 literature labeled strains, of which 33 are tailocin producers. Results are given in Table 2.

#### Case study (PTE and prophage distribution across soil bacteria and plant-associated bacteria isolated from different parts of plants) and statistical analysis

We applied PhageTAILor to 7,925 bacterial genomes from the IMG database, partitioned into 4 isolation niches by their metadata, namely Soil (n = 4,018), root-associated (n = 2,511), plant-associated (n = 846), and leaf-associated (n = 550). Genomes were first downloaded with the following criteria: (Taxonomy -- Domain [Bacteria]) AND (Sequencing Assembly Annotation -- Sequencing Status [Finished, Permanent Draft]) AND (Environmental Classification -- Ecosystem [Environmental, Host-associated]) AND (Environmental Classification -- Ecosystem Category [Plants, Terrestrial]) AND (Genome Statistics Metadata -- Scaffold Count (Number of scaffolds) (Range: 0 to 633334083) [1 to 100]). Among these genomes, manual curation was done to exclude rock surface samples, heavily contaminated pasture soil samples with animal manure or blood, beach sand, cave samples, meteorite samples, bog and wetland samples, high salinity soil samples, geothermal soil samples, anthropogenically contaminated soil samples, ancient soil preserved in amber samples, industrial site soil samples, extraterrestrial site samples, landfill soil samples, compost soil samples, aquifer samples, deep subsurface samples, and plant aboveground part samples. Then samples isolated from leaves, phylloplane or phyllosphere were classified as leaf-associated samples, samples isolated from plant root, rhizosphere or rhizoplane were classified as root-associated samples, and samples isolated from plant without description on isolation from root or leaf were classified as plant-associated samples. Soil samples lacking plant association were classified as soil samples. These niche classification and taxonomy of these resulting 7,925 bacterial genomes are shown in Figure 3.

To avoid re-running geNomad across this dataset, provirus regions and predictions were extracted from a prior Microbial Virus Pipeline run on the identical genome set. The remaining five detectors were executed de novo on Pyrodigal-gv predicted proteins to populate all downstream feature blocks. For each PTE class and niche contrast, we constructed 2 x 2 contingency tables and evaluated significance using a two-sided Fisher’s exact test (scipy.stats.fisher_exact, SciPy v1.11) (53). Pooled comparisons aggregated the three plant-associated niches into a single plant stratum tested against bulk soil, while per-niche comparisons evaluated each plant niche individually against bulk soil, with odds ratios expressed relative to soil. To confirm that niche contrasts persist after accounting for taxonomic composition, we calculated taxonomy-adjusted odds ratios for each plant niche and the pooled plant stratum relative to bulk soil. Adjustments were made using the Mantel–Haenszel estimator stratified by GTDB order, with 95% confidence intervals calculated via the Robins–Breslow–Greenland method and conditional association evaluated using the Mantel–Haenszel chi-square test. Taxa with zero genomes in a given stratum contributed zero weight to the corresponding estimate. The twelve per niche adjusted tests are interpreted against a Bonferroni threshold of 0.05 divided by 12.

Our niche level results both recapitulate and refine earlier large-scale isolate genome comparisons of these systems, while also revealing informative contrasts. Previous surveys showed that eCIS loci are enriched in environmental microbes from soil, terrestrial, and aquatic habitats, but depleted in mammalian and avian pathogens (3). Similarly, comparative analyses of isolate genomes revealed that T6SSs are enriched in plant-associated bacteria and depleted in bulk soil, whereas eCIS elements are abundant in both environments (25). At the unadjusted level, our genome-scale survey aligns with these observations: T6SS and tailocin prevalence increases from bulk soil to plant-associated niches, while eCIS loci exhibit a soil-biased distribution. Crucially, our dataset decouples environmental habitat from taxonomic lineage. After stratifying by GTDB order using a Mantel–Haenszel estimator, the pooled plant-versus-soil enrichment of T6SS collapsed. This indicates that much of the apparent plant signal is compositional, reflecting the taxonomic orders that colonize plants rather than a true niche -level adaptation—though a modest rhizosphere-specific T6SS enrichment and a phyllosphere reduction persisted. In contrast, the soil bias of eCIS remained robust after taxonomic adjustment, confirming a consistent depletion among plant-associated bacteria.This is consistent with the earlier picture of eCIS as an environmental, soil common system, and it adds the resolution that within environmental microbes eCIS leans toward soil over the plant niches rather than being evenly spread across them. Taken together, these comparisons place PhageTAILor’s calls in line with prior biology and supply the taxonomy-controlled resolution needed to distinguish a true niche association from one driven by sampling.

### Part B. User pipeline and outputs

The user pipeline accepts a tab separated sample sheet with a stable sample number, a unique sample name, and a path to each assembly, and is invoked through the bundled phagetailor command, which wraps Snakemake and provides plain language per step documentation. The pipeline executes ten stages in order, namely gene calling with Pyrodigal-gv, taxonomy assignment with GTDB-Tk plus a skani search against the bundled training sketch, five parallel candidate detectors, an HMM search, feature extraction, classification with the 4 heads, and report generation. The workflow is modular, so any single detector can be run on its own, and precomputed results such as an existing geNomad run can be reused by staging them at the expected output paths. Requesting classification runs all six detectors, which keeps the user inference feature distribution identical to training. The pipeline produces four primary outputs, a per-region prediction table with all four head probabilities and the confidence band, a per-genome summary with the GTDB-Tk taxonomy and ANI band, a BED file of all candidate regions for genome browsers such as IGV, JBrowse, and the UCSC Genome Browser (54–56), and a self contained browser readable report, together with one raw candidate table per detector preserved as a first class output (Figure 1C).

## Data availability

PhageTAILor is available on Github (https://github.com/hjcho-bio/PhageTAILor). Data files are available on Zenodo (10.5281/zenodo.21152308).

## ACKNOWLEDGEMENTS

This work was supported by the Lawrence Berkeley National Laboratory Directed Research and Development Program (LBNL LDRD #23-105). The US Department of Energy (DOE) Joint Genome Institute, a DOE Office of Science user facility, is supported by the Office of Science of the US Department of Energy under Contract No. DE-AC02-05CH11231. We thank Maureen Berg for helpful feedback and comments on this manuscript. We appreciate Britt Koskella and Dominique Holtappels for sharing their phyllosphere data. We appreciate Catherine Mageeney Ellis Torrance, Hannah McClain and Kelly Williams for their help on TIGER software. We also thank Jennifer Yuzon for providing foundation input on several components of this pipeline. This material is, in part, based upon work supported by the National Science Foundation Biology Integration Institute grant (NSF Award No. 2119968; PI-Ceballos), which support S.H. and R.M.C. Any opinions, findings and conclusions or recommendations expressed in this material are those of the authors and do not necessarily reflect the views of the National Science Foundation. Socheata Hour was supported by the US Department of Energy (DOE) RENEW grant No. DE-SC0024237.

## AUTHOR CONTRIBUTIONS

J.T., R.B. and S.R. conceived and designed the study with input from authors. H.C conducted pipeline development. H.C., S.H., T.S. performed bioinformatic analyses, which were run on the informatics cluster of the Joint Genome Institute, Lawrence Berkeley National Laboratory. S.R., C.C., O.A. and L.A. provided contributions to experimental design, methods and data interpretation. J.T., R.B. and H.C. drafted the manuscript with contributions from authors. Authors read and approved the final paper.

## ETHICS DECLARATIONS

Claude was used to optimize the human-generated scripts, Claude and Gemini were used for grammar correction of human-generated texts and logical integrity check, Perplexity was used to assist literature search, and Paperplot was used to get better ideas on diagrams. All generative AI tool outputs and suggestions were validated and curated by human authors.

## DATASETS

**Figure S1.**
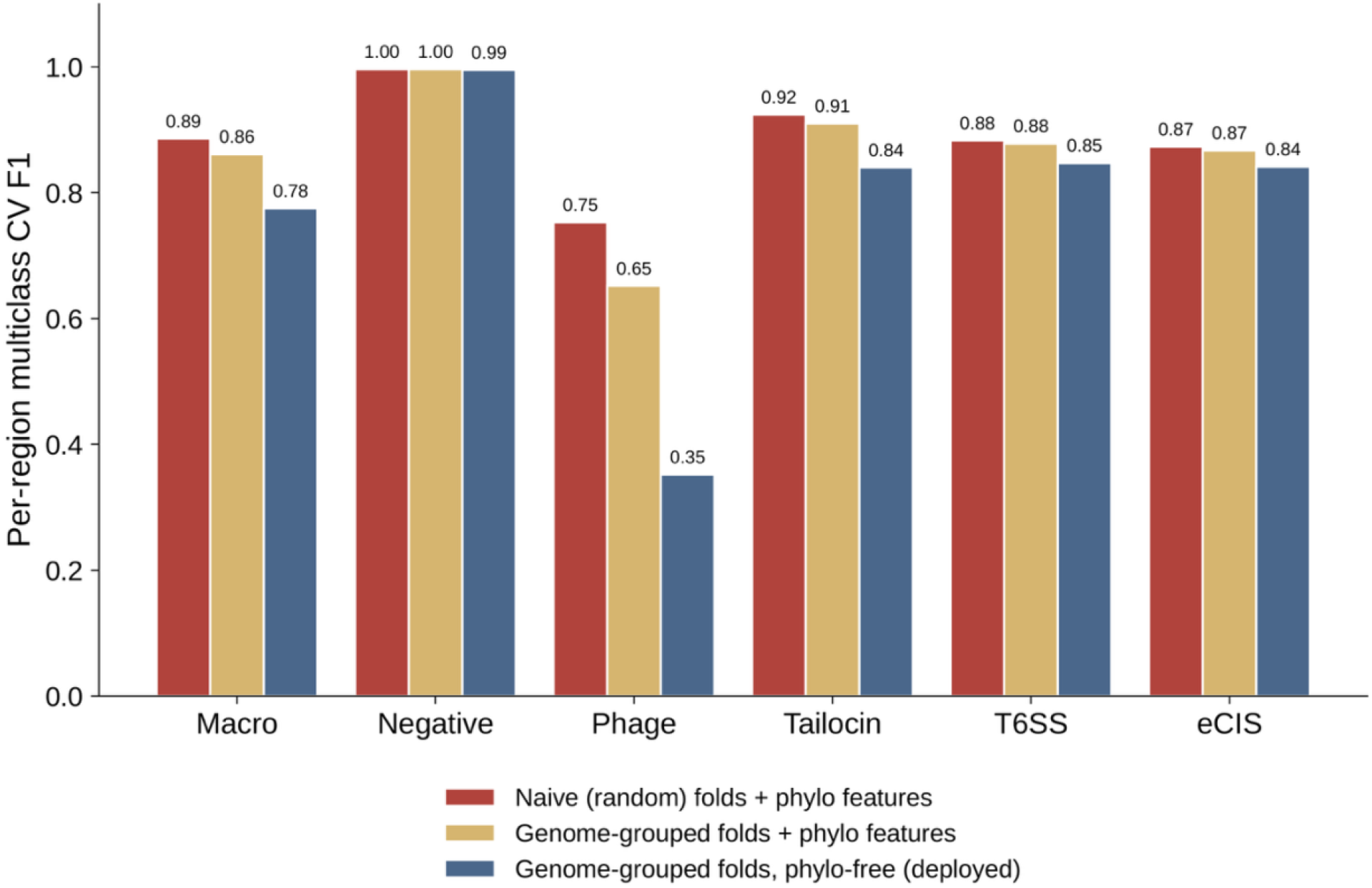
Cross-validation leakage decomposition. Macro and per-class F1 of the per-region multiclass model under three evaluation settings that progressively remove sources of optimism: (1) naive random folds with the five phylogenetic-placement features; (2) genome-grouped folds with those features; (3) genome-grouped folds without them (the deployed phylo-free model). The modest decline (macro F1 0.886 → 0.861 → 0.775) shows that within-genome leakage and genus memorization are both small.

**Figure S2.**
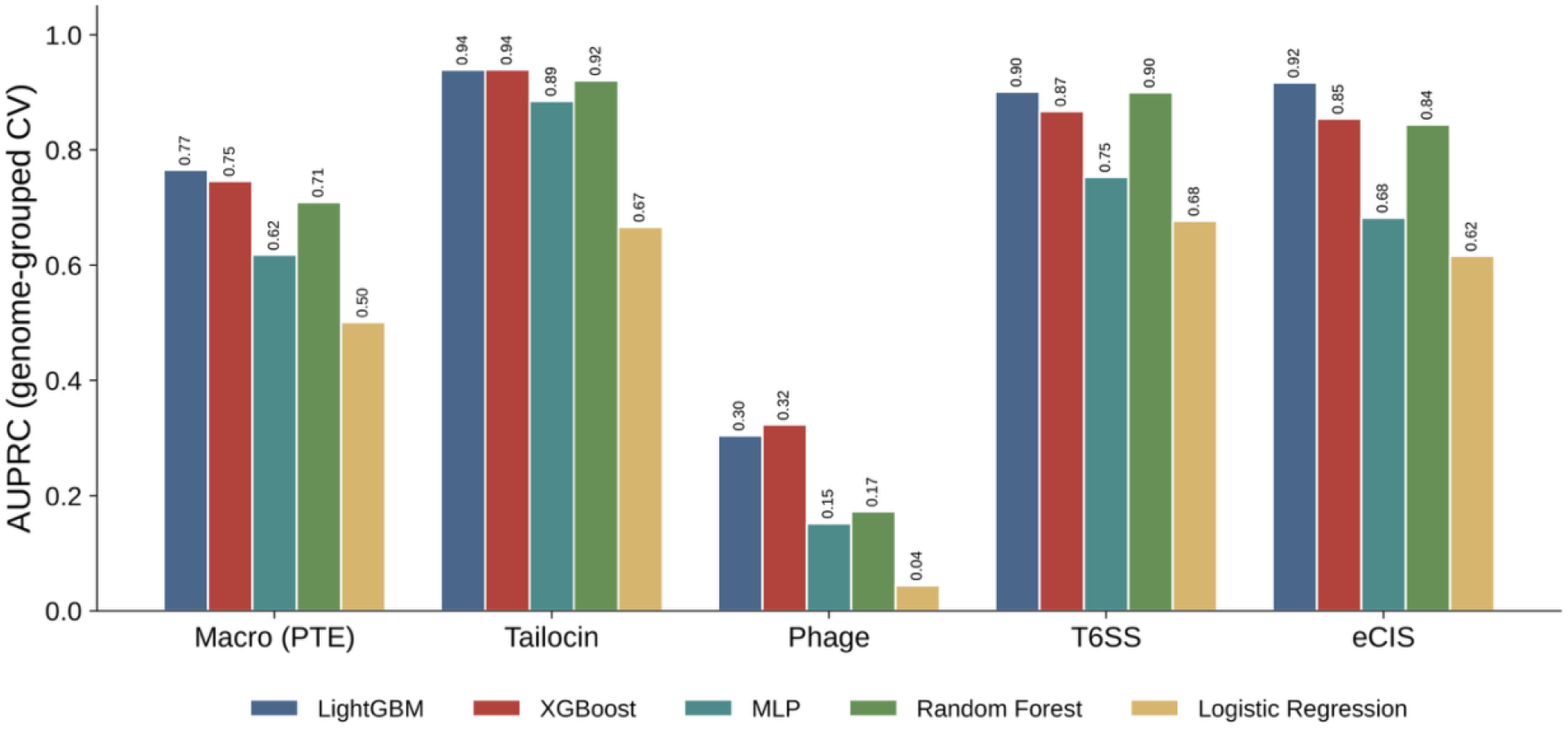
Algorithm comparison by per-class AUPRC. Five classifiers (LightGBM, XGBoost, MLP, Random Forest, logistic regression) trained on the identical phylo-free 74-feature matrix under genome-grouped 5-fold cross-validation (GroupKFold by genome); per-class one-vs-rest AUPRC computed from pooled out-of-fold probabilities — a threshold-free, weighting-independent ranking metric that avoids the instability of argmax macro-F1 under extreme class imbalance. LightGBM is selected for deployment.

**Figure S3.**
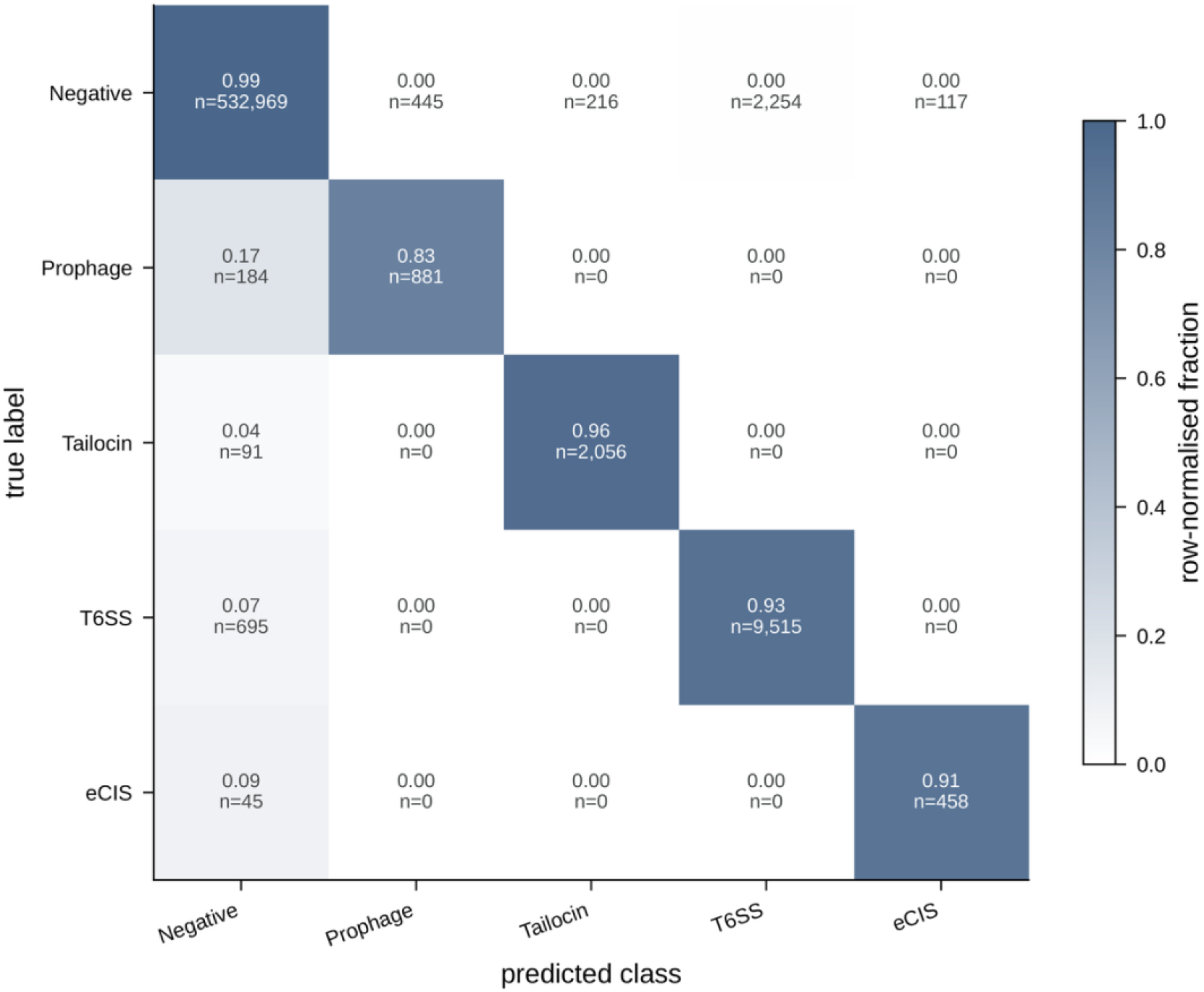
Five-class confusion matrix. Out-of-fold predictions of the multiclass head under 5-fold cross-validation, as a 5 × 5 matrix (rows, true label; columns, predicted label; negative, prophage, tailocin, T6SS, eCIS), row-normalized to the fraction of each true class assigned to each predicted class (cell color = row-normalized fraction; counts annotated).

**Figure S4.**
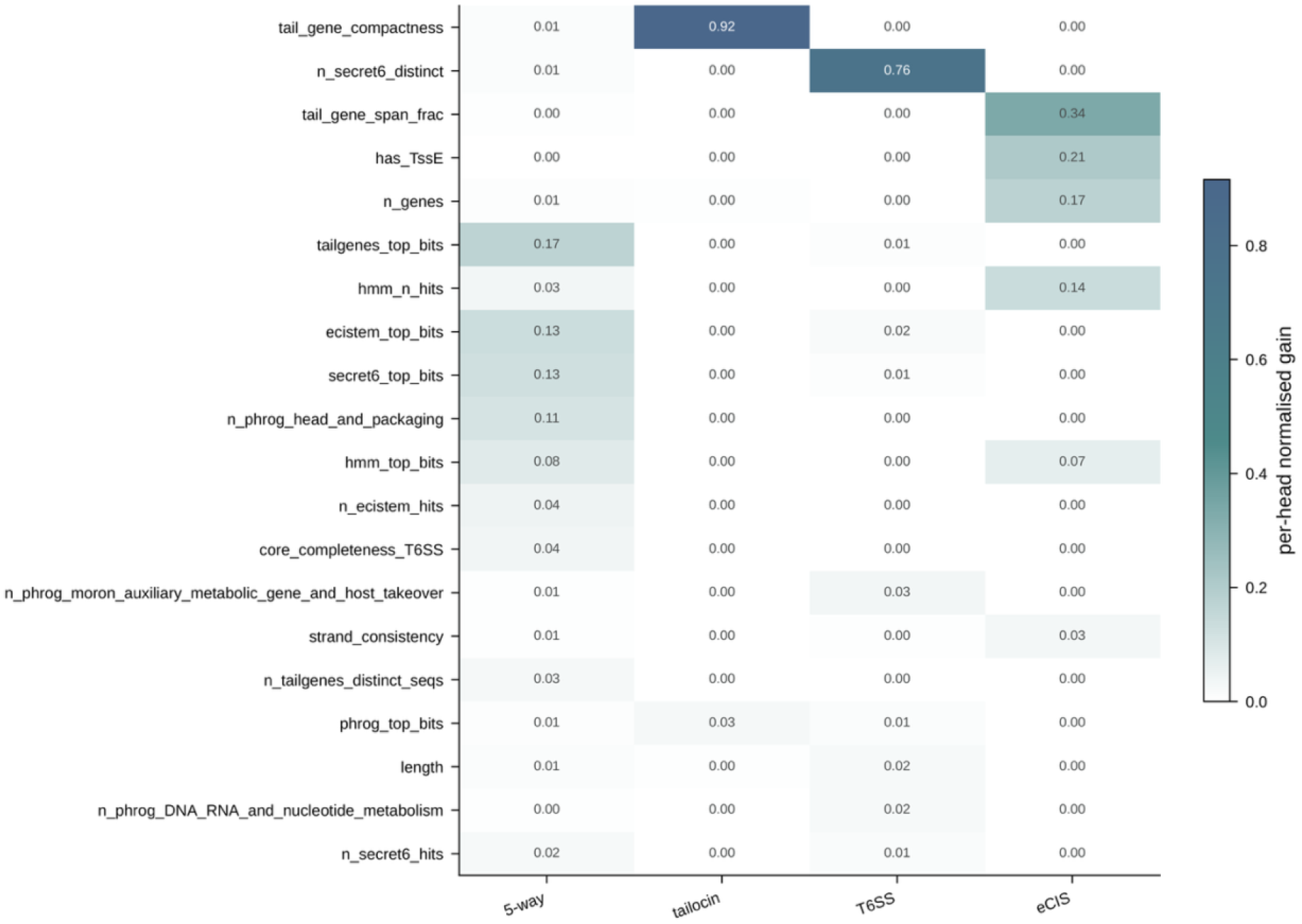
Top feature importances across the four model heads. Heatmap of the top 20 features by LightGBM gain for each of the four boosters (the 5-way multiclass head and the tailocin, T6SS, and eCIS binary heads); gain is normalized within each head so the heads are comparable (color = per-head normalized gain).

**Figure S5.**
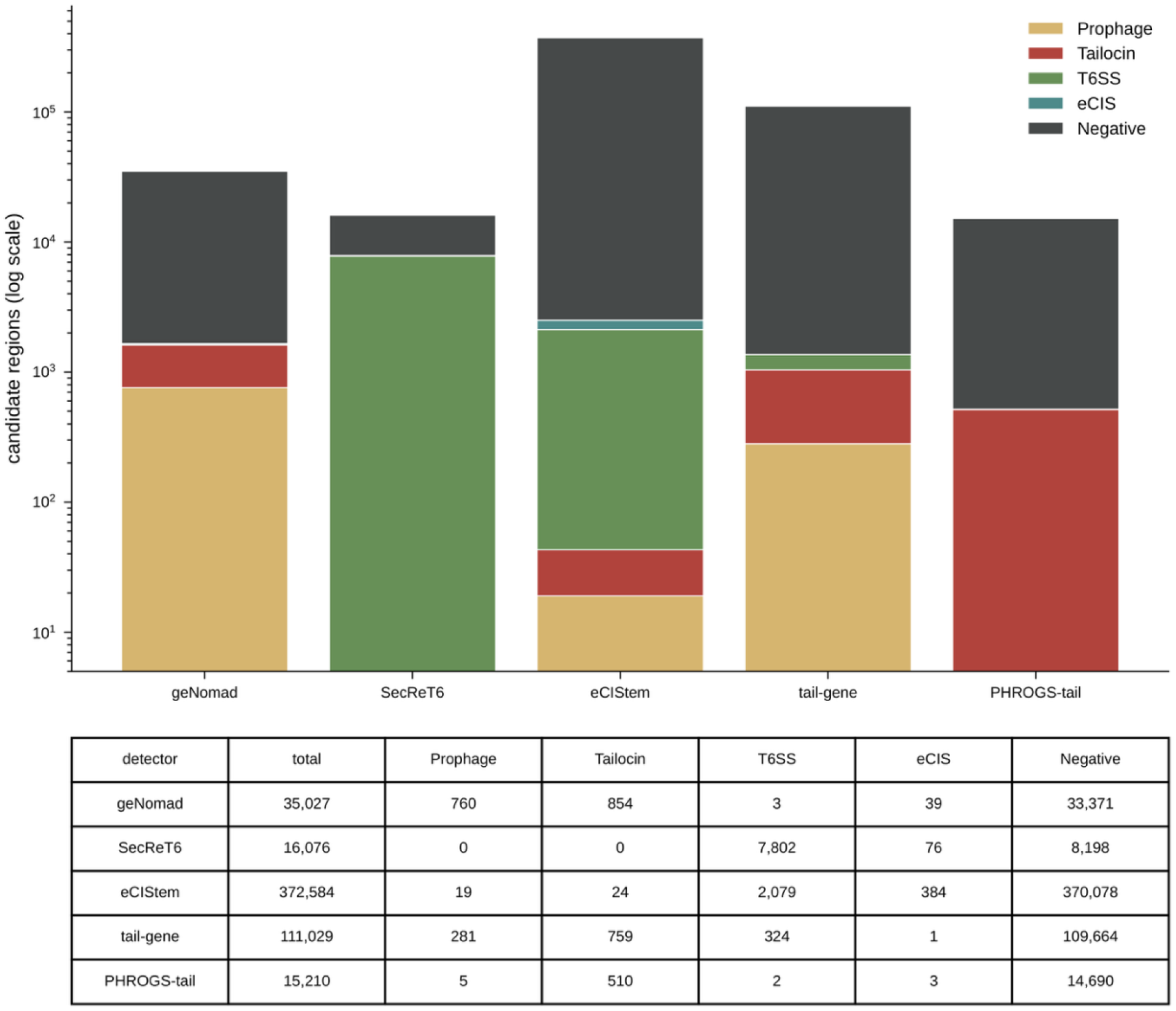
Per-detector candidate counts, stacked by final class label. Number of candidate regions contributed by each of the six detectors (geNomad, tail-gene, PHROGS-tail, SecReT6, eCIStem, tail-HMM), stacked by the region’s final predicted class, with a per-detector × class table beneath the bars. A standalone, expanded view of the detector breakdown summarized in the pipeline figure.

**Figure S6.**
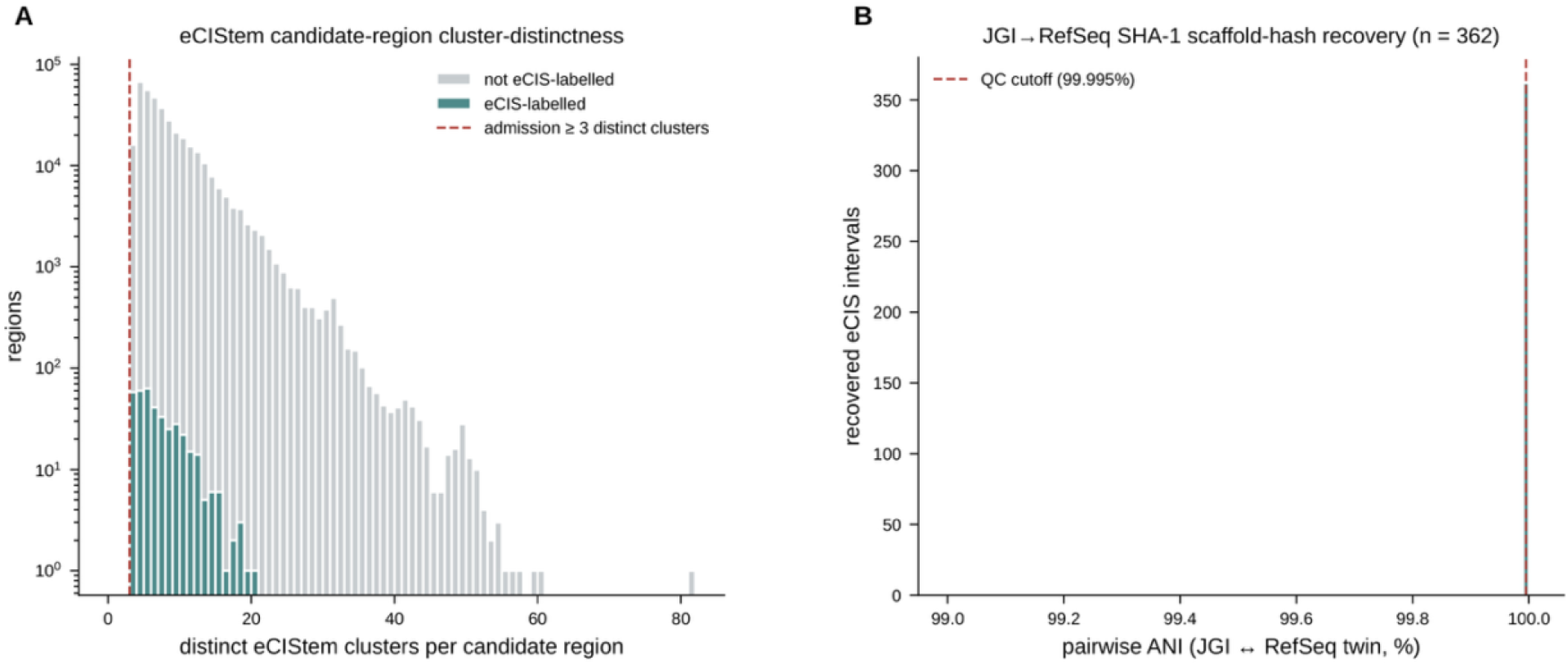
eCIS detection deep-dive. **(A)** Distribution of the number of distinct eCIStem clusters matched per eCIS candidate region, with the ≥ 3-cluster admission threshold marked and the label-positive eCIS fraction annotated. **(B)** ANI distribution of the 477 JGI eCIS intervals recovered to their RefSeq training twins by SHA-1 scaffold-hash matching, relative to those twins.

**Figure S7.**
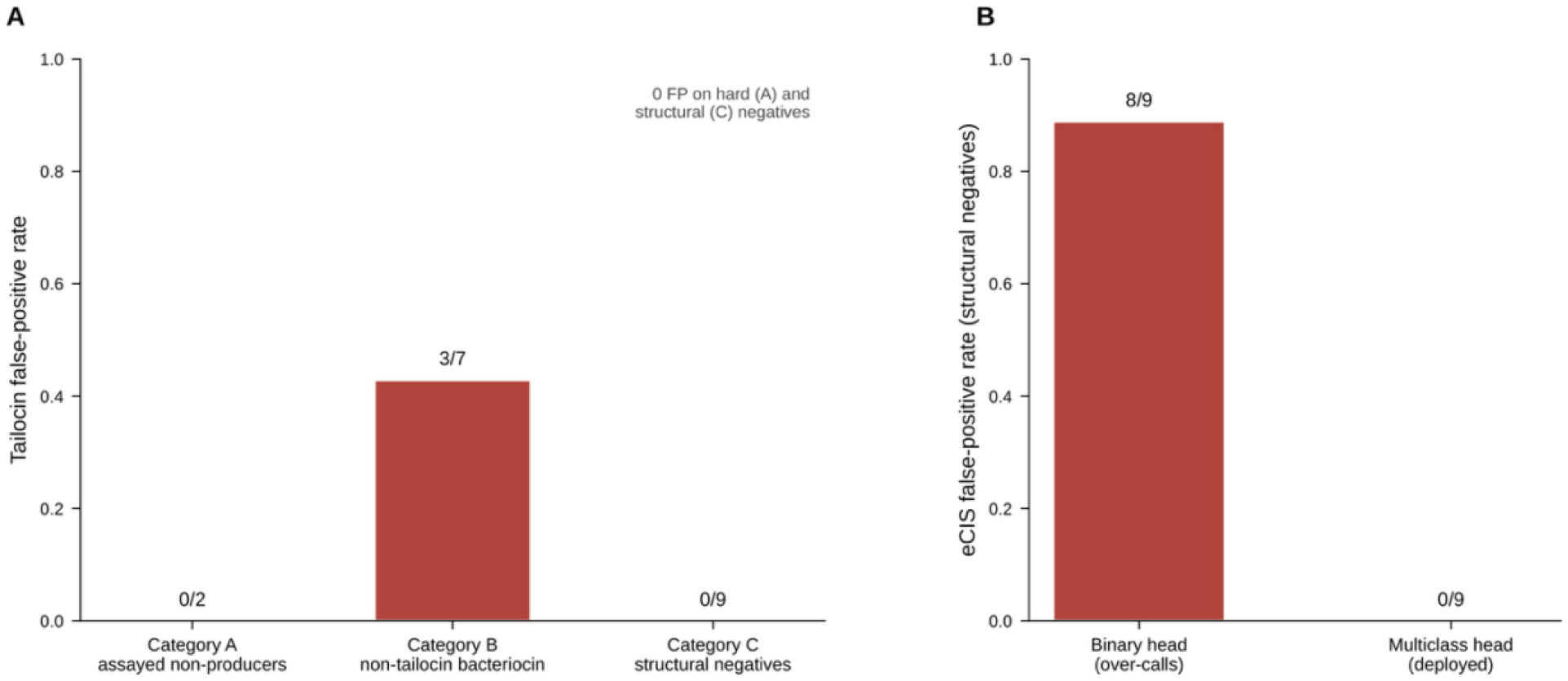
Clean-negative precision panel. False-positive behavior on a curated panel of genomes that credibly lack tailocins, across 3 negative categories: Category A, experimentally assayed non-producers (hard, same-clade); Category B, non-contractile bacteriocin producers with no reported tailocin; Category C, obligate-intracellular or minimal genomes structurally lacking contractile machinery. **(A)** Tailocin false-positive rate by category — zero on the hard category A and structural category C, with false positives only on prophage-bearing category B genomes (overall tailocin precision 0.81 at recall 1.00). **(B)** eCIS false-positive rate on the category C structural negatives: the binary eCIS head over-calls (8/9), whereas the deployed multiclass eCIS head scores them ≈ 10⁻⁸ (0/9) — the reason eCIS calls are taken from the multiclass head.

**Figure S8.**
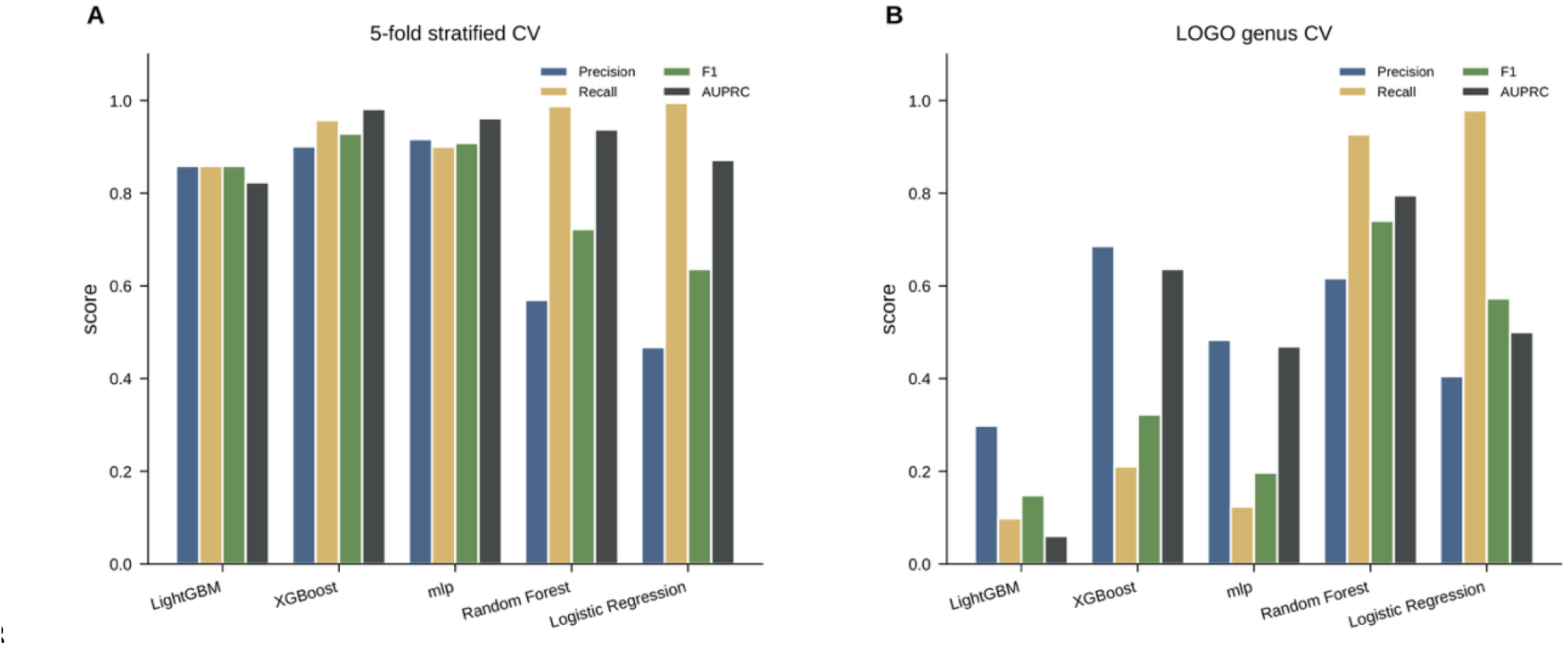
Algorithm comparison on the binary tailocin task. Performance of five classifiers on the binary tailocin task from algorithm_comparison.tsv, including leave-one-genus-out generalization. XGBoost and LightGBM are essentially tied, Random Forest fails to generalize under leave-one-genus-out, and logistic regression is unsuitable. This is the conservative, fully supported binary counterpart to the multiclass comparison in Fig. S2.

**Figure S9.**
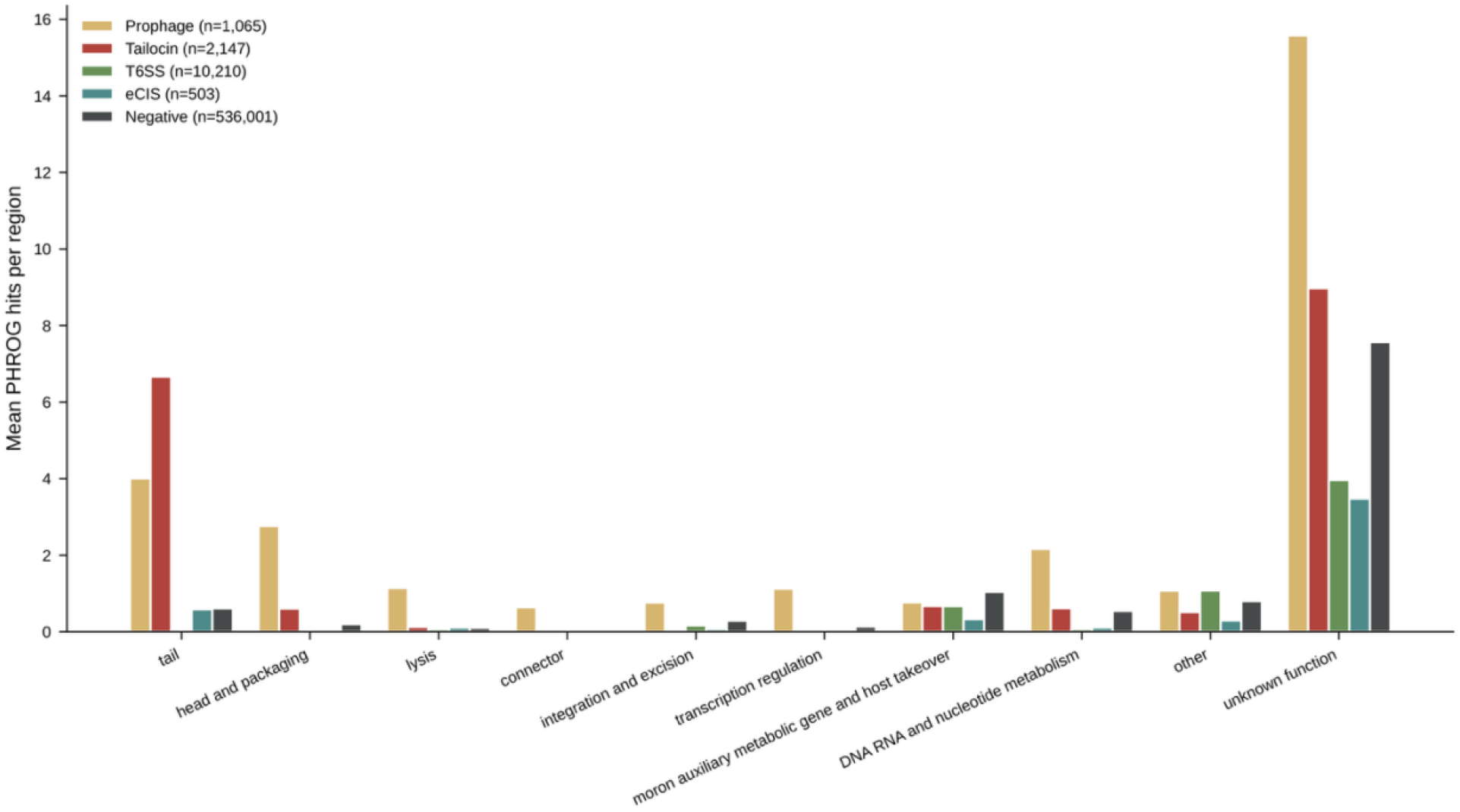
PHROG functional-category content per candidate region, by class. Mean number of PHROG hits per candidate region in each PHROG functional category (tail; head and packaging; lysis; connector; integration and excision; transcription regulation; moron, auxiliary metabolic gene and host takeover; DNA, RNA and nucleotide metabolism; other; unknown function), grouped by region class (prophage, tailocin, T6SS, eCIS, and negative; per-class n shown). Tail-category content is the dominant discriminator of contractile elements.

**Figure S10.**
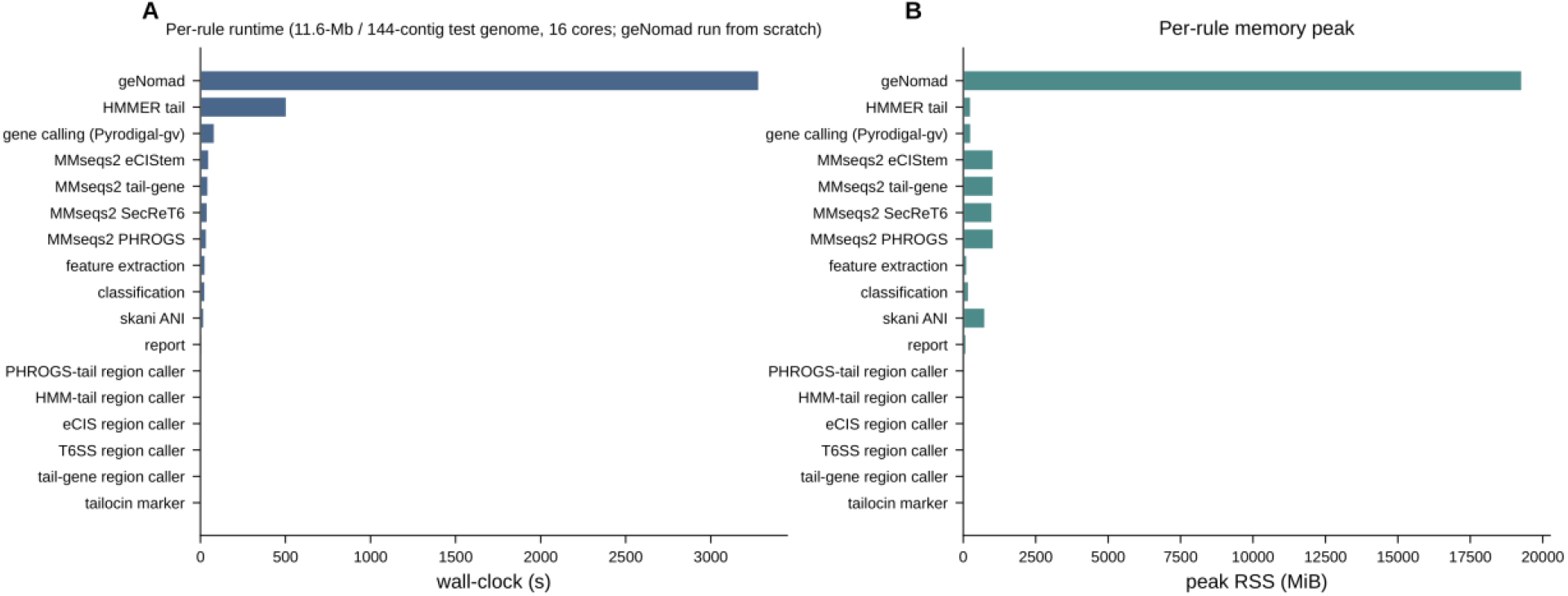
Pipeline runtime and memory by step. Wall-clock time and peak resident memory for each pipeline step (gene calling, taxonomy, geNomad, the four MMseqs2 detectors, tail-HMM/feature assembly, classification, and report) on a representative 5-Mb genome.

**Figure S11.**
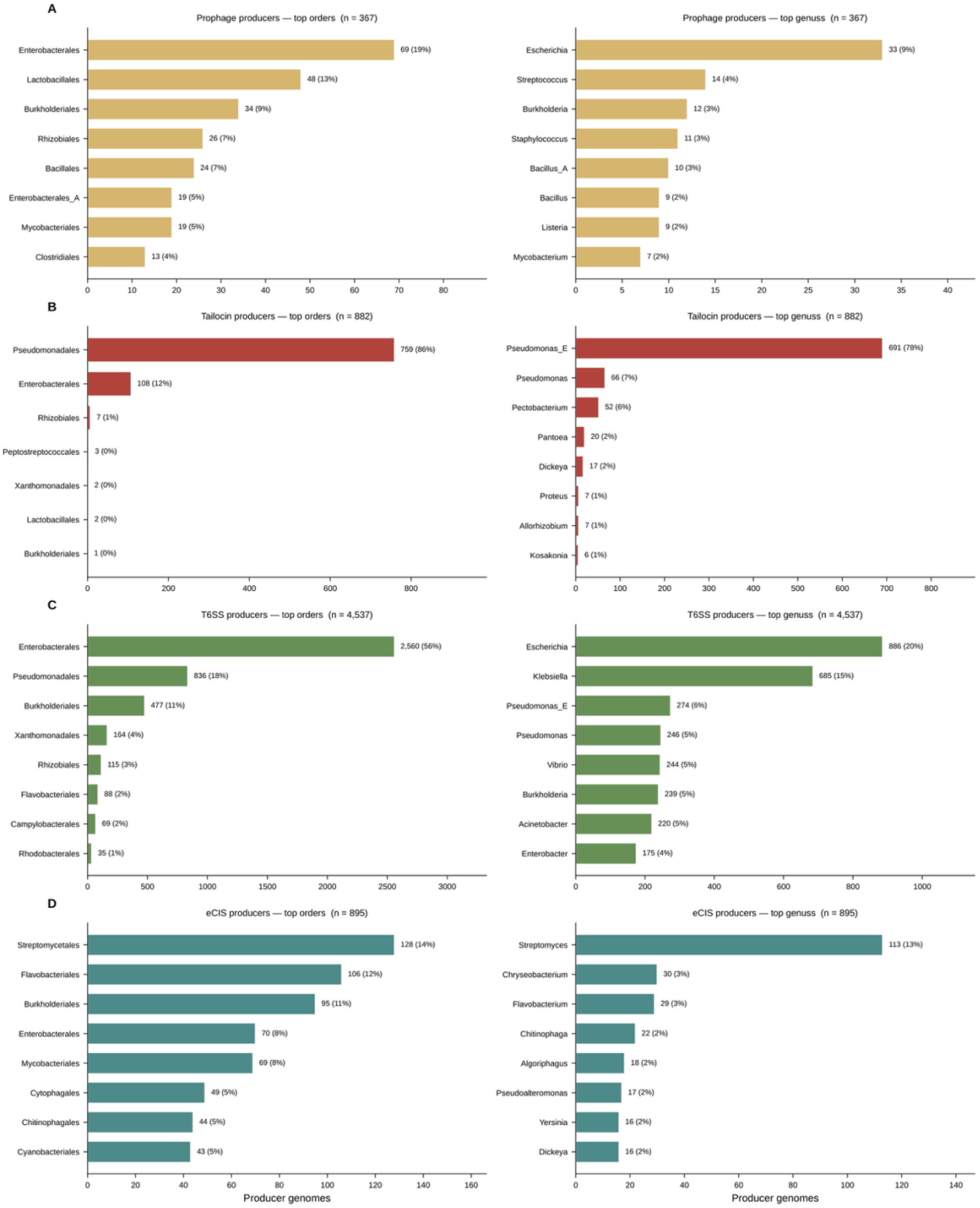
Producer taxonomy by PTE class. Producer-genome counts by GTDB order (left column) and genus (right column), one row per prophage or PTE class (tailocin, prophage, T6SS, eCIS); horizontal bars show the top eight taxa per cell as a percentage of that class’s producers. The per-class lineage skew hidden by a phylum-level view is evident — for example, tailocin producers are overwhelmingly *Pseudomonas*, whereas T6SS producers are dominated by Enterobacterales.

**Figure S12.**
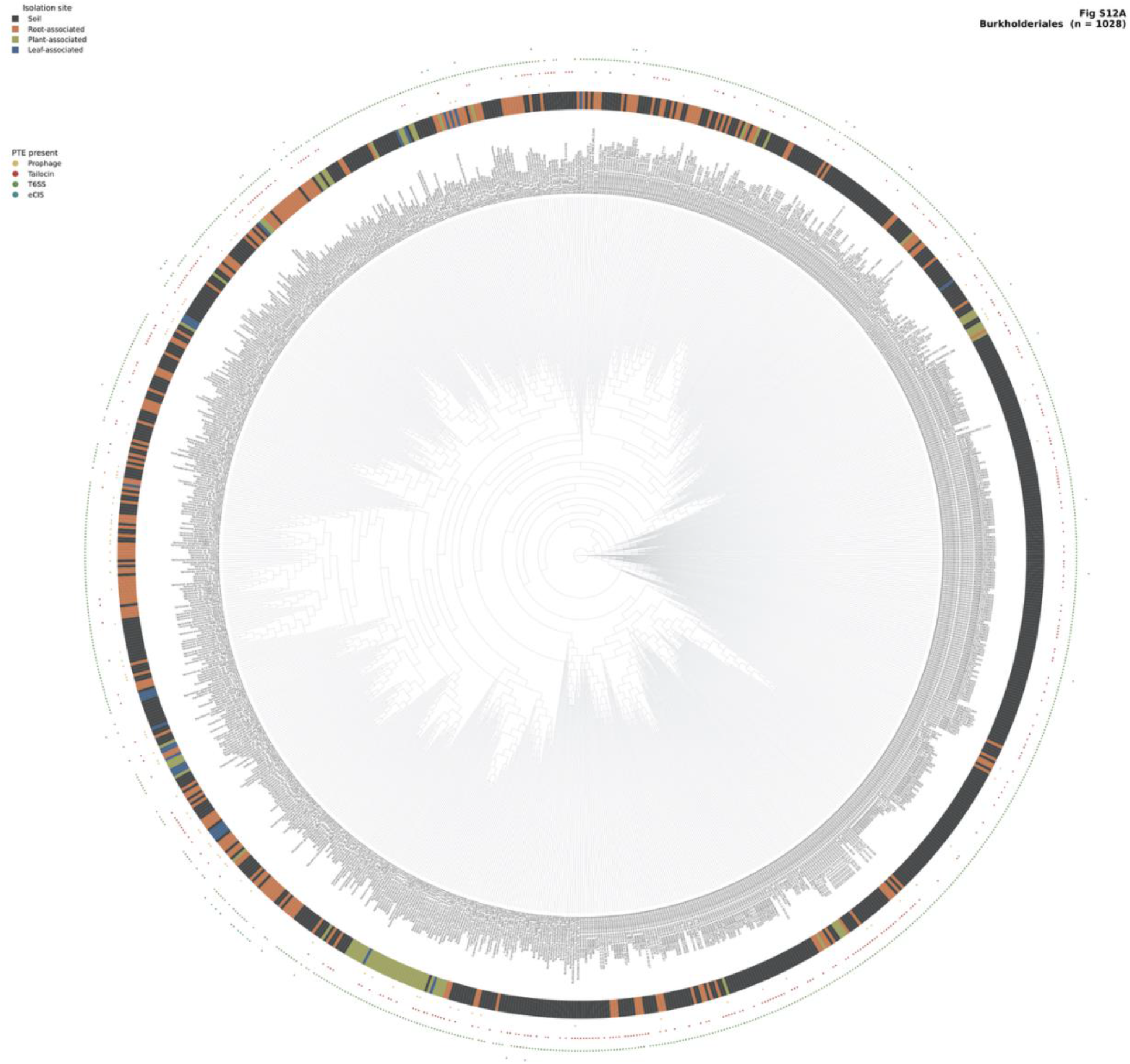
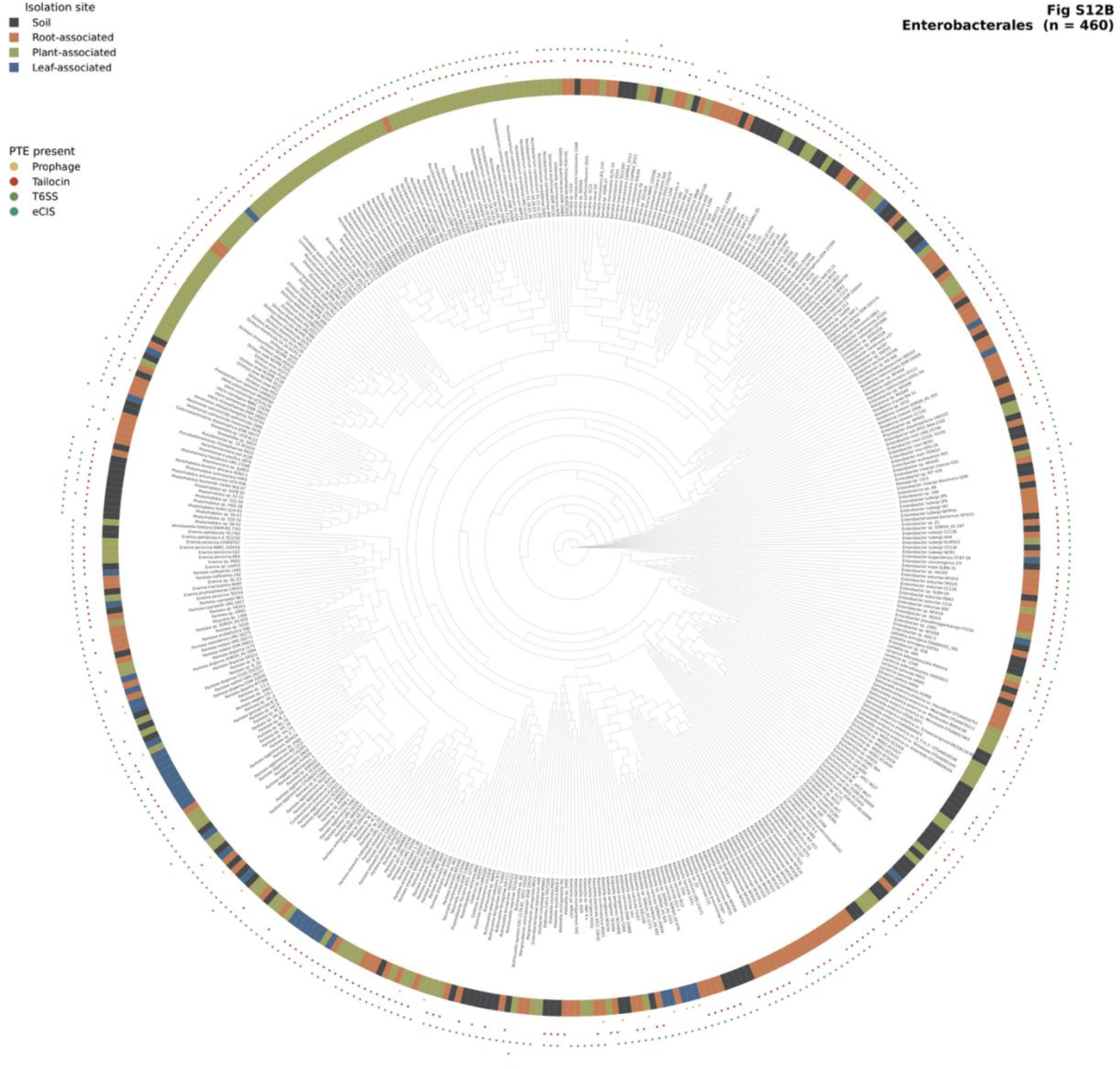
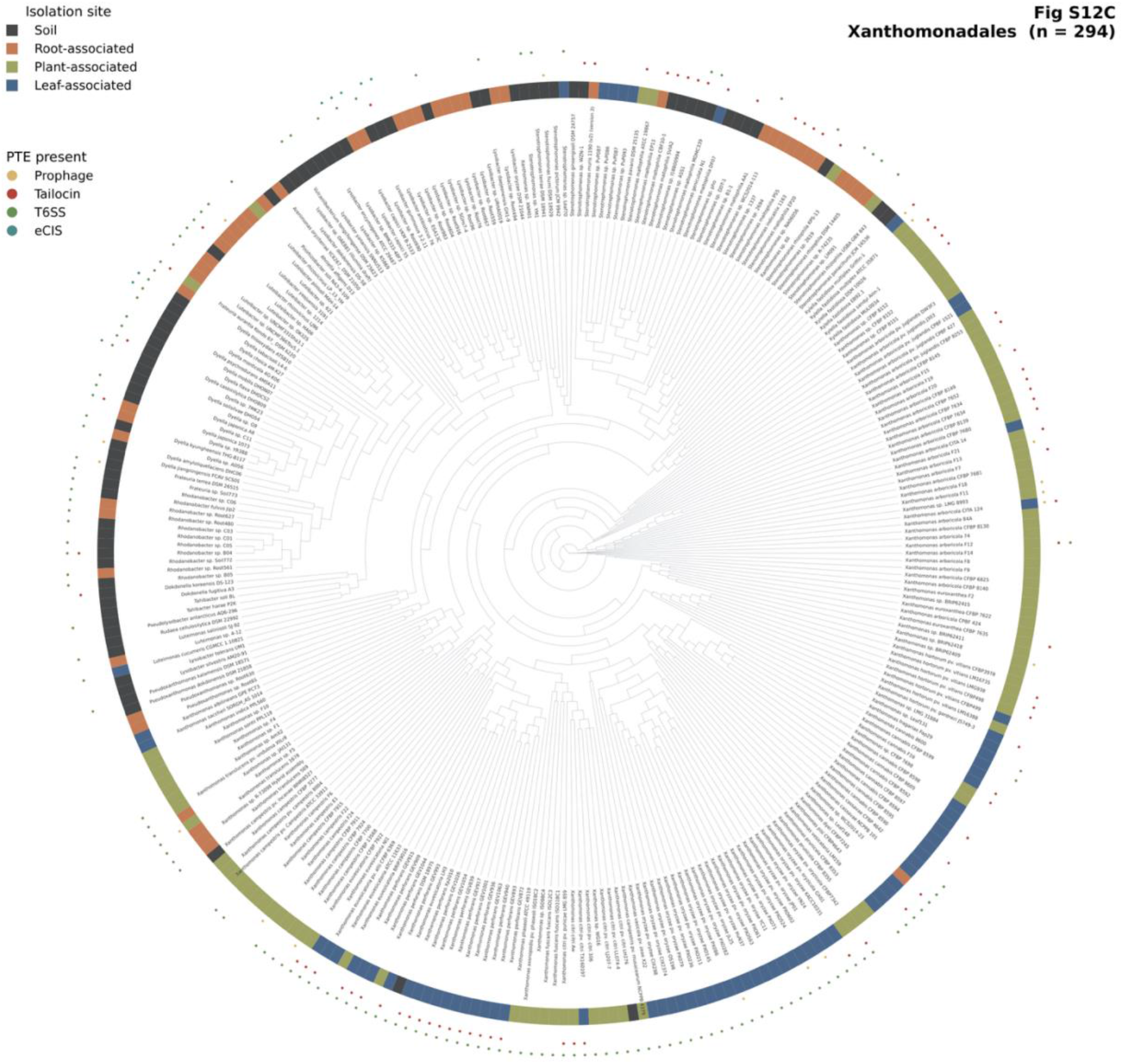
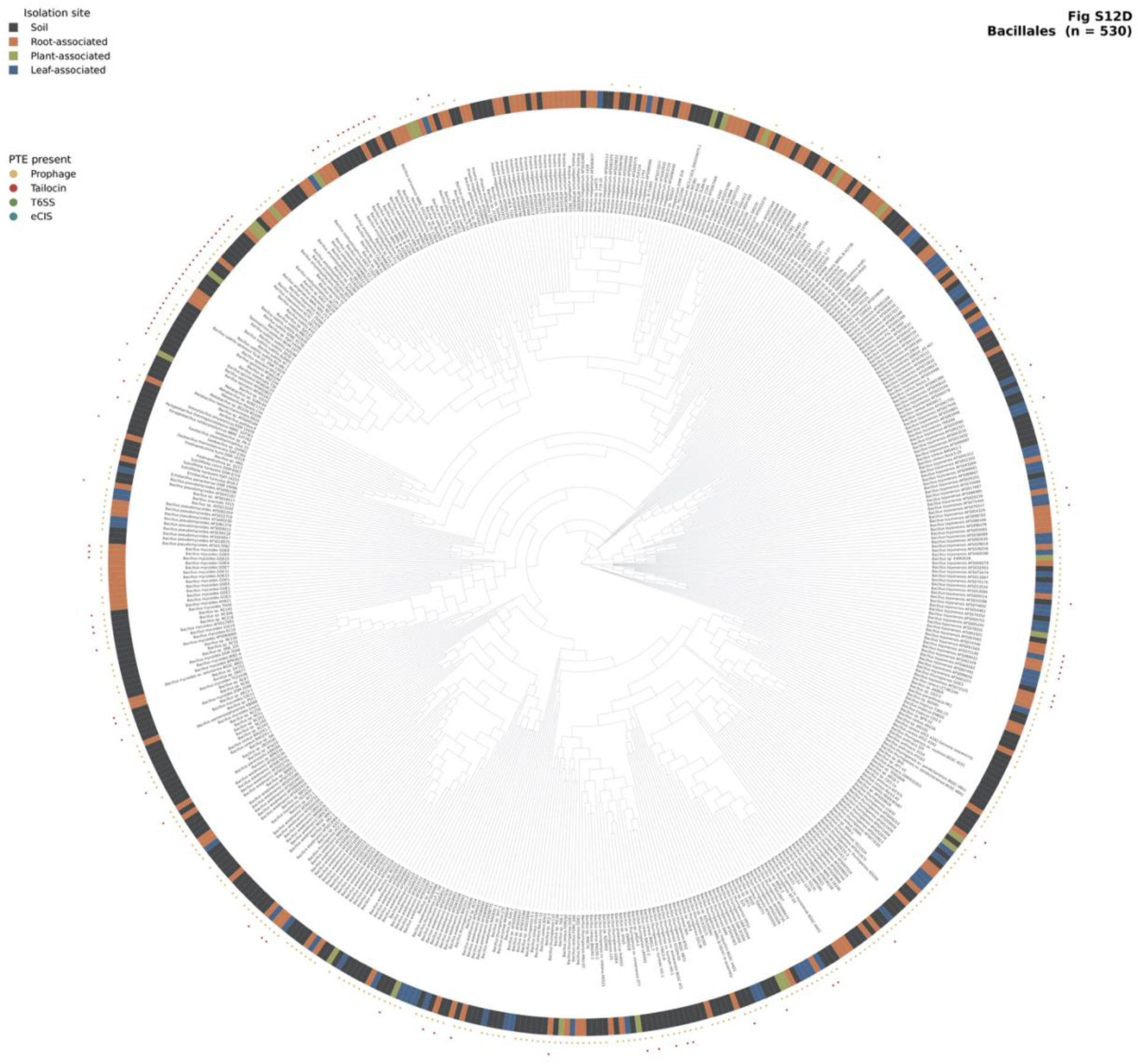
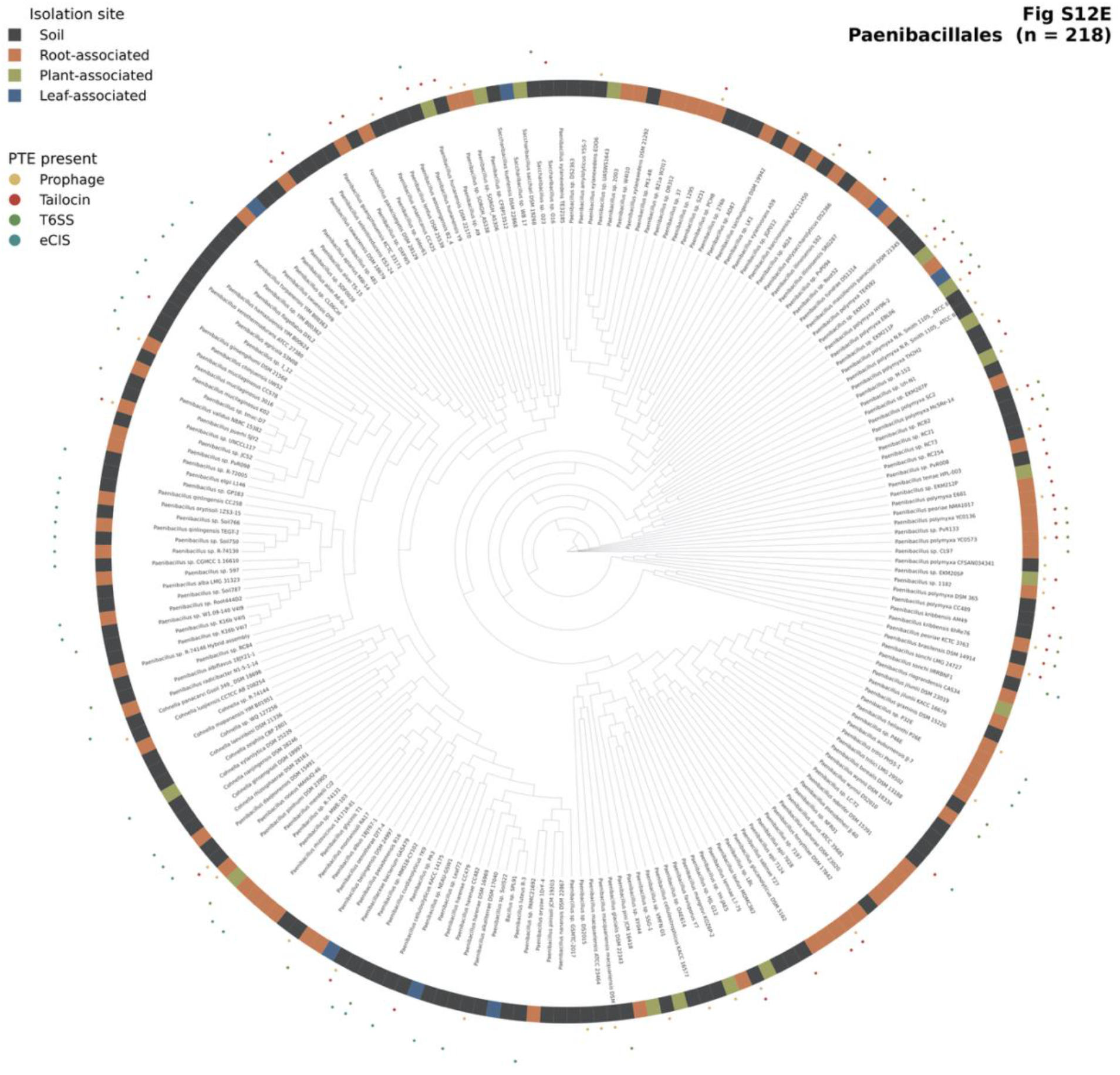
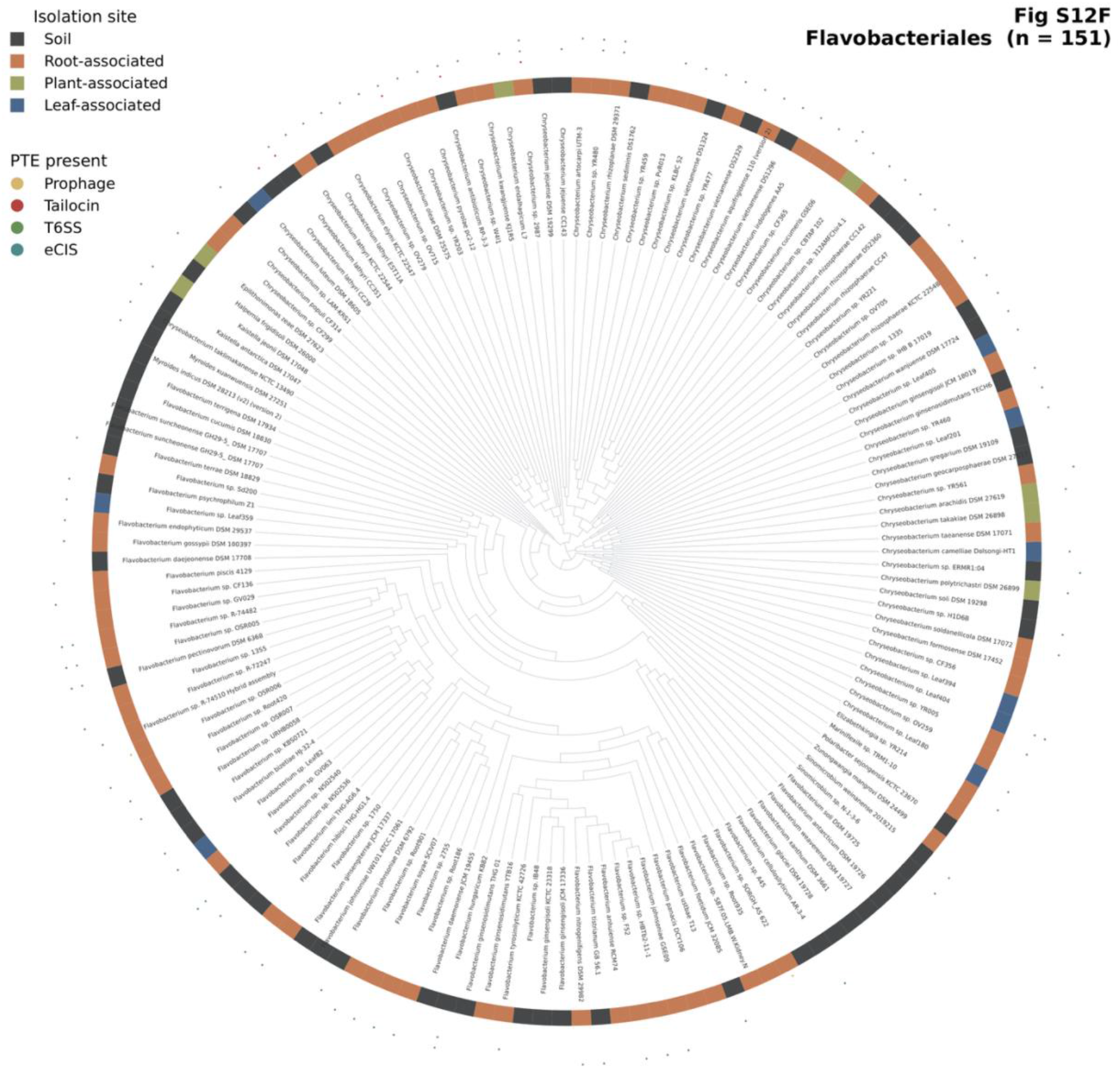
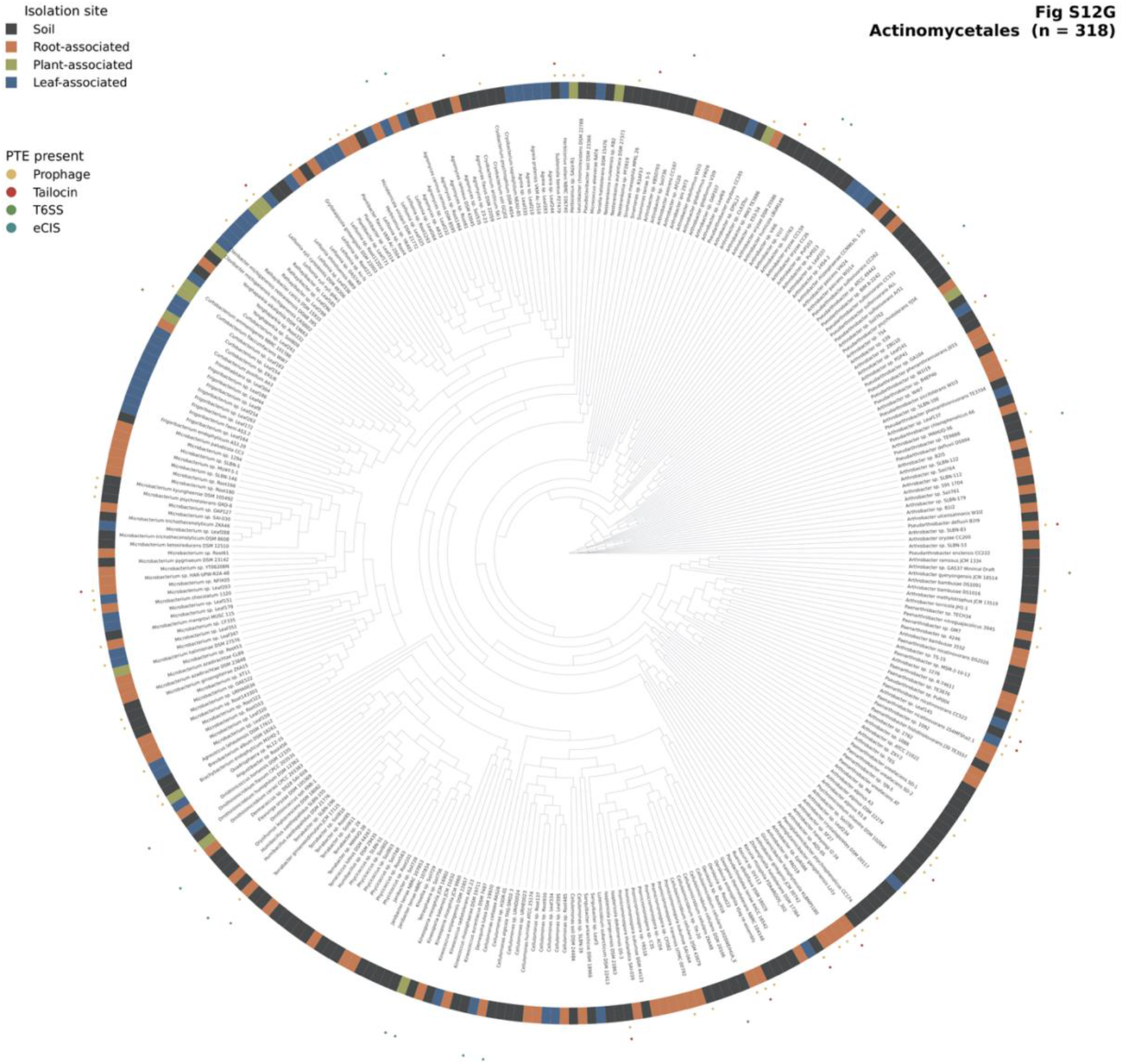
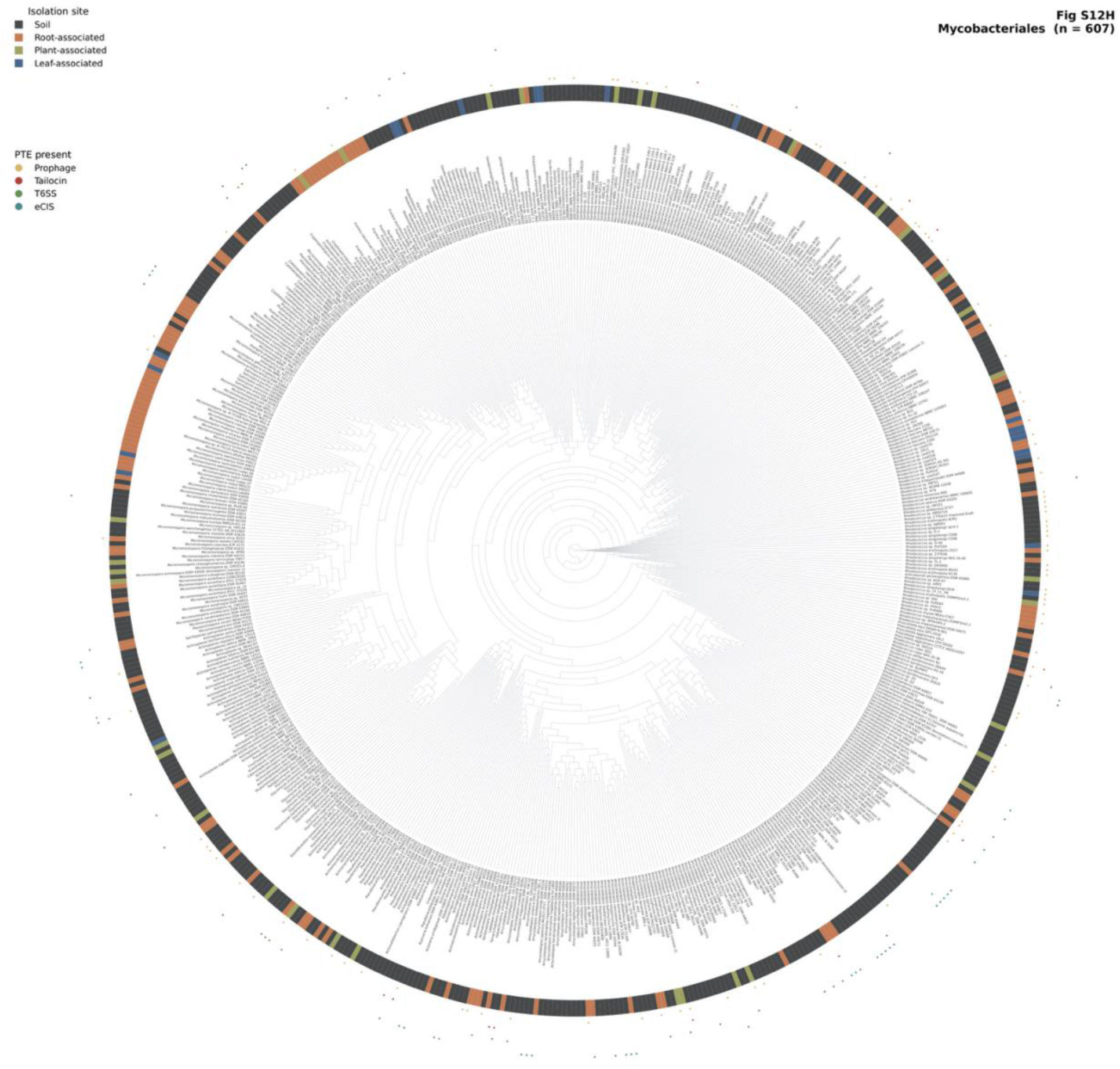
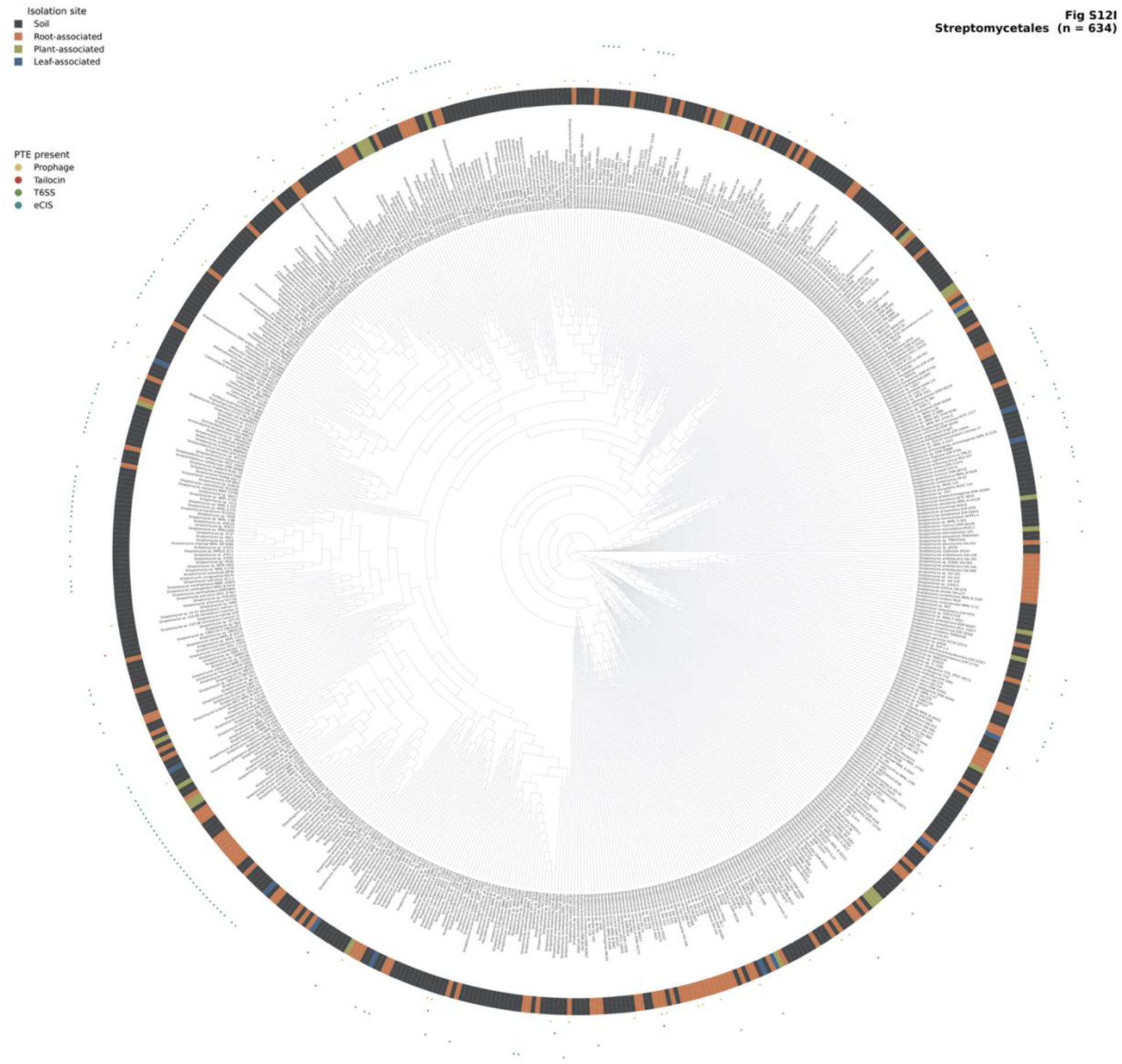
Per-order phylogenetic distribution of prophages and PTEs for the remaining 9 orders. Circular cladograms for the nine abundant orders from the 7,925-genome plant- and soil-associated cohort not shown in Fig. 6, grouped by phylum (top, Pseudomonadota: Burkholderiales, Enterobacterales, Lysobacterales; middle, Bacillota and Bacteroidota: Bacillales, Paenibacillales, Flavobacteriales; bottom, Actinomycetota: Actinomycetales, Mycobacteriales, Kitasatosporales). Inner ring, isolation niche; four outer dot rings, prophage or PTE-class presence (deployed calls); per-tip names omitted. Across orders, prophage or PTE carriage is phylogenetically structured and does not co-distribute with niche.

---

**Supplementary Table 1.** Composition and per-strain calls of the Tier 4 tailocin evaluation panel: 13 experimentally validated cross-genus producers and 18 curated tailocin-free negatives (categories A/B/C), with the deployed-model and genus-held-out calls.

**Supplementary Table 2.** A. Producer-genome list: 6,776 prophages and PTE records over 6,501 unique genomes, with genome size, scaffold count, CheckM2 QC, source DOI, and prophage or PTE class. B. All 13,082-training prophage and PTE coordinate intervals (scaffold, start, end, type) with the per-row coordinate-source DOI.

**Supplementary Table 3.** A. JGI IMG metadata of 7,925 genomes used for the case study, comparing prophage and PTE distribution across plant-associated, root-associated, leaf-associated, and soil-associated bacteria. B. Per-genome prophage or PTE calls (tailocin, prophage, T6SS, eCIS) with GTDB order and isolation niche for the 7,925-genome set. C. Per-niche crude (Fisher) and taxonomy-adjusted (Mantel-Haenszel, stratified by GTDB order) odds ratios with 95% CIs per prophage or PTE class (Figure 3F).

**Supplementary Table 4.** Per-strain predictions of the PhageTAILor model on the held-out cross-clade benchmark (per-class probabilities, binary-head scores, ANI band, expected label).

**Supplementary Table 5.** PhageTAILor model head-to-head tool comparison on the held-out benchmark (PhageTAILor v0.1.0 phylo-free, TattleTail, geNomad-standalone).

**Supplementary Table 6.** Annotated 74-feature per-region column list used by all four deployed heads (the phylo-free matrix; phylogenetic-placement features excluded).

**Supplementary Table 7.** Full software inventory (channel, dependency, pinned version) parsed from workflow/envs/*.yaml.

**Supplementary Table 8.** Statistics of the 7,925 plant-, root-, leaf-associated and soil bacteria.

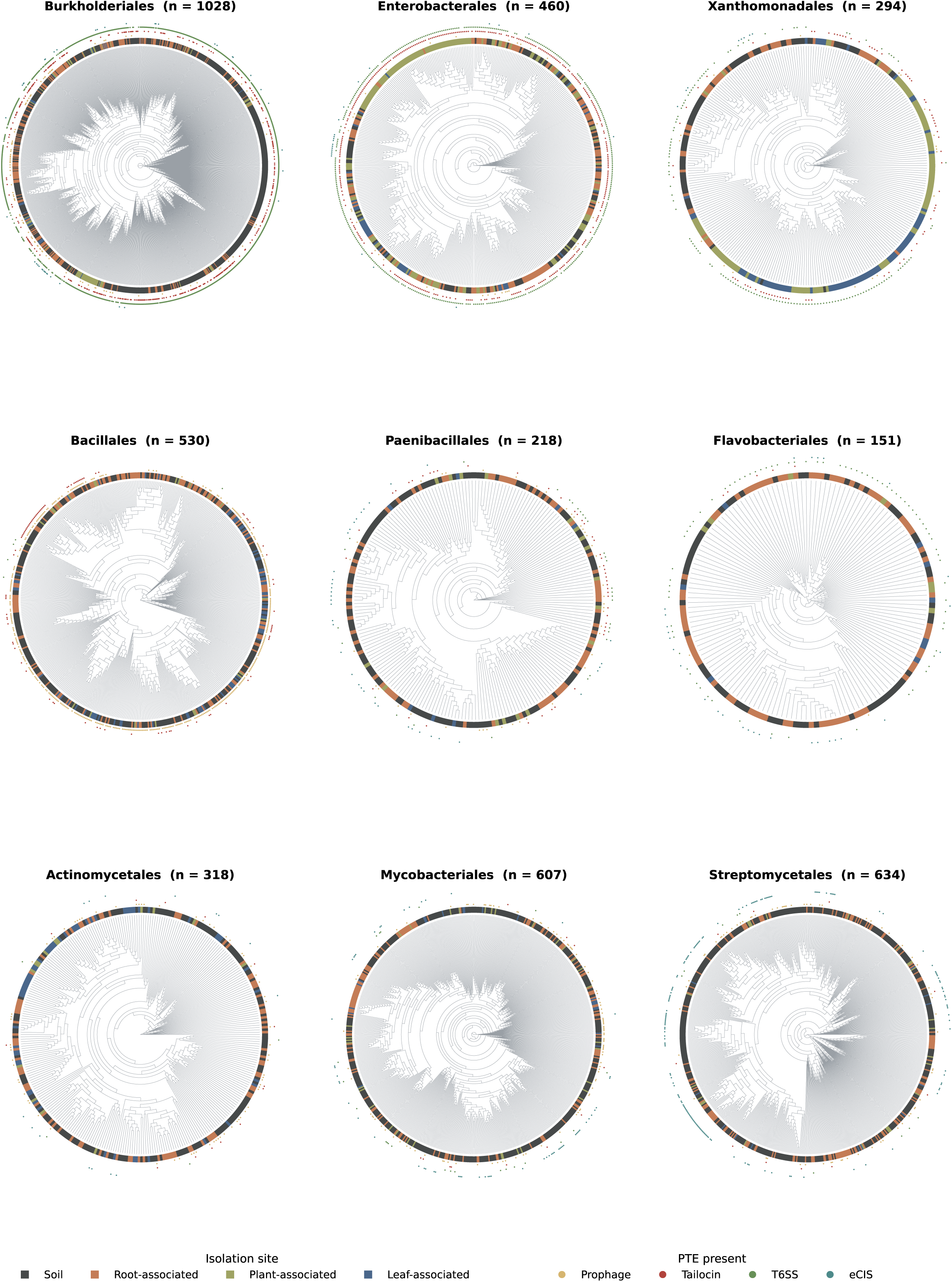

